# High-throughput glioblastoma-neurovascular assembloids model blood-tumor barrier disruption, tumor reprogramming and therapeutic efficacy

**DOI:** 10.1101/2024.11.11.622979

**Authors:** Pei Zhuang, Shuai Gao, Benjamin Scott, Yi-Bo Jia, Wei-Min Meng, Rui Yin, Xin-Yu Nie, Soniya Tamhankar, Sean P. Palecek, Ludovica Gaiaschi, Jorge L. Jimenez Macias, Shuichi Mizuno, Sean E. Lawler, Eric V. Shusta, Martine Lamfers, Feng-Feng Bei, Choi-Fong Cho

## Abstract

The blood-tumor barrier (BTB) in glioblastoma is an abnormal neurovascular interface that can be focally leaky yet still restrict effective therapy. We present a rapid human BTB model in which patient-derived glioblastoma stem cells (GSCs), human brain endothelial cells, pericytes and astrocytes self-assemble within 72 hours into uniform three-dimensional “BTB assembloids”. Thousands can be generated in parallel in standard plates, enabling high-throughput studies and drug testing. These assembloids establish spatial glioma-vascular and glioma-astrocyte niches with focal loss of endothelial tight-junction proteins, astrocyte recruitment and endothelial recruitment toward tumor clusters with localized junctional disruption at the tumor-vascular interface. BTB-conditioned GSCs acquire more invasive, stemness, mesenchymal and angiogenic programs compared with matched GSC neurospheres, and these BTB-conditioned states translate into greater engraftment and infiltrative growth *in vivo,* accompanied by vascular remodeling and BTB-like barrier disruption. The assembloid platform supports quantitative, ‘barrier-aware’ assessment of compound access and retention in human BTB- (and BBB)-like contexts, revealing that functional P-glycoprotein efflux persists despite increased BTB permeability, thereby dissociating structural barrier leakiness from therapeutic accessibility. It is modular and compatible with iPSC-derived brain cells (for example brain endothelial cells), microglia or other immune components, thereby providing a route toward genotype- and immunity-specific modeling. Together, this establishes BTB assembloids as a scalable human system to interrogate glioblastoma-neurovascular crosstalk and to evaluate brain-tumor therapeutics.

---

Malignant gliomas, particularly glioblastoma (GBM), remain uniformly lethal despite maximal resection and concurrent chemoradiotherapy, and responses to immunotherapies have been modest^1, 2^. A major stumbling block for effective therapeutic development for brain tumors is the presence of the blood-brain barrier (BBB) that hinders drug delivery to tumor sites^3^. The BBB features a unique and intricate structural arrangement of the neurovascular unit (NVU), primarily composed of tightly connected endothelial cells (EC) that are closely associated with astrocytes, pericytes, neurons and other cell types, which are critical for maintaining barrier integrity^4^. In brain tumors, the BBB is partially preserved and pathologically remodeled within tumors into a heterogeneous blood-tumor barrier (BTB), producing niches that are variably permeable yet often therapeutically refractory^5^. The BTB arises from complex and highly dynamic interaction and mechanisms between the tumor cells and surrounding NVU, leading to focal tight-junction loss, altered transport/efflux and perivascular remodeling that together shape invasion and treatment response^6,7^.

Preclinical neuro-oncology research heavily depends on animal models, which are costly, low throughput, slow to establish (some tumors taking months to establish), and limited by interspecies differences in both tumor biology and barrier function^8^. *In vitro* models offer control and scalability but have notable gaps. Culture models including cancer cell lines and glioma stem cell (GSC) neurospheres are often used as a starting point, providing valuable information to complement *in vivo* findings^9^. To date, relatively few studies include tumor cells in BBB culture models^7^. Most widely described *in vitro* BTB systems involve the 2D transwell model, where brain EC (along with other NVU cells) are typically cultured on a semiporous membrane on the upper apical side, while GBM cells are cultured in the lower basal chamber^10,11^, allowing exposure of the cells to the conditioned medium without being in direct contact. Despite its prevalence owing to easy accessibility and versatility, the transwell model has several well-known limitations including the inability to faithfully capture the complex 3D architecture of the BTB and direct tumor-NVU interactions which are known to be critical for tumor plasticity. Microfluidic “BTB-on-a-chip” platforms introduce flow and lumenized endothelium but require specialized hardware and are difficult to scale for discovery-stage screening^12^.

Brain organoids represent a new frontier for modeling the complex physiology of the human brain and neurological diseases^13,14^. GBM organoids (GBO) derived directly from patient tumors capture intratumoral heterogeneity and clinically relevant histopathology, and can be biobanked for longitudinal studies^8,15,16^. Most protocols reach experimental maturity over ∼2-4 weeks (sometimes longer)^8,17–19^, which is feasible for precision oncology timelines but can constrain early discovery throughput and iteration cadence^20^. Reported challenges include batch-to-batch variability, size/composition heterogeneity within and across organoids, as well as protocol-dependent differences that complicate cross-study comparisons^21,22^. Recent benchmarking and rigor frameworks are helping, but variability remains a recognized issue for scalable screening. Moreover, while vascularization strategies (genetic induction of endothelial programs or fusion with vessel organoids) can impart BBB-like features^23–26^, these approaches add complexity and often reduce scalability relative to plate-based assays. Patient-derived GBOs have been successfully engrafted into mouse brain, where they exhibit robust vascularization, growth and histopathology that mirror patient disease^27–29^; however, the new vessels are largely host-derived (mouse) rather than human. This chimerism is powerful for tumor biology in a native brain milieu but limits “human BTB” transport fidelity because endothelial junctions, transporters and efflux programs are species-specific.

We have previously described a human BBB assembloid model that reproduces key BBB marker expression and barrier functions, as well as predicts BBB permeability *in vivo*^30–37^. Here, we incorporate patient-derived GSC to rapidly form human “BTB assembloids” that reconstruct essential components of the glioma-neurovascular unit. Within 3 days, uniform three-dimensional BTB assembloids self-organize at scale (hundreds-thousands per run in standard multi-well plates), enabling parallel, large-scale testing of conditions and quantitative evaluation of compound permeability and tumor retention across the BTB (and BBB). The assembloids establish spatially organized glioma-endothelial and glioma-astrocyte niches with focal tight-junction disruption, astrocyte recruitment and abnormal vascular-like structures that resemble clinical tumor microenvironments. Comparative transcriptomic and functional analyses show that GSCs within the BTB assembloids rapidly upregulate stem-like, invasive, mesenchymal, and pro-angiogenic pathways relative to their matched GSC neurospheres (NS) alone. This BTB-conditioned state increases engraftment and infiltrative growth *in vivo*. In parallel, endothelial/perivascular compartments adopt BTB-like phenotypes. This system is modular, whereby induced pluripotent stem cell (iPSC)-derived brain cells, microglia or other immune cells, and patient-matched components can be incorporated to interrogate genotype- and immunity-specific mechanisms.

We further demonstrate the utility of the BTB assembloid for modeling therapeutic delivery to GBM cells. This platform operationalizes ‘barrier-aware’ therapeutic evaluation by quantifying uptake/retention and efficacy of a well-validated GBM-targeting peptide (e.g. BTP-7)^33,36^, brain-penetrant vector (AAV.CPP.16)^31^ and standard chemotherapies (e.g. temozolomide and doxorubicin) across the BTB, linking delivery to tumor response. These studies highlight the proof-of-concept translational utility of the BTB assembloid platform for assessing drug permeability and optimizing therapeutic strategies against GBM.

## Results

### Establishment of the BTB assembloids

BTB assembloids were generated through co-culturing patient-derived GSC neurospheres, human brain endothelial cells (EC), human brain vascular pericytes (HBVP) and human astrocytes (HA), which self-organized into a multicellular BTB microtissue. Immunofluorescence staining of cell-specific biomarkers confirmed the identity of the HBVP through PDGFR-β expression, HA through glial fibrillary acidic protein (GFAP) expression, and hCMEC through CD31 and CD105 expression (**Extended Data Fig. 1a**)^30,35^.

GSCs are normally maintained in the absence of serum in Neurobasal (NB) media to preserve stem-like properties^9^. Prolonged culture of GSCs in 10% FBS can erode stemness and drive divergence from the parental tumor profile^9,38^. Our BBB assembloids are cultured in media containing 1.5% human serum (HS)^30^, so we first assessed whether this low-serum condition is compatible with GSC stemness in a mixed NVU co-culture.

The BTB assembloids were generated by a two-step self-assembly (**Fig. 1a**): **1)** GSCs were aggregated for 24 h in serum-free NB to form small neurospheres; **2)** HA, HBVP and cerebral microvascular endothelial cells (hCMEC) were co-cultured with the neurospheres for 48 h in either 1.5% HS or NB. We applied this to 5 GSC models (G9, G30, GBM-X6, GS184, GS401) - three of which are derived from patients (GBM-X6, GS184, GS401), with GS184 (primary) and GS401 (recurrent) paired from the same patient. Each line displayed distinct BTB phenotypes (**Extended Data Fig. 1b**). For example, under HS condition, GS184 formed compact tumor clumps, whereas GS401 was more infiltrative and dispersed. In all cases, GSC neurospheres initially embedded within the assembloid, then began to proliferate and invade toward the periphery over time. BTB assembloids reached ∼300-400 μm in diameter (vs 150-250 μm for BBB assembloids) and remained viable for several weeks.

**Fig. 1.**
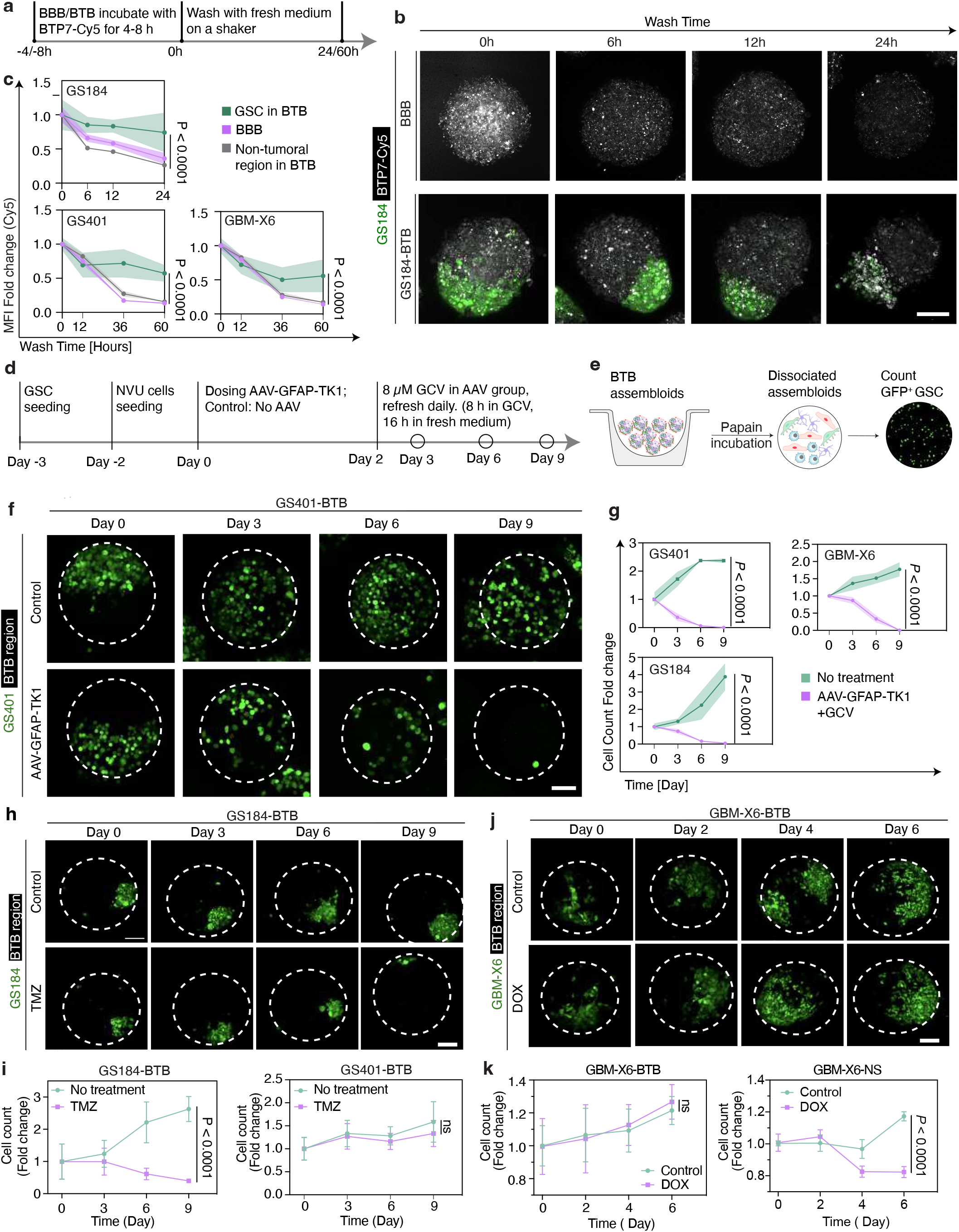
Blood-tumor barrier (BTB) assembloid formation and structural organization. **(a)** The BTB assembloid was established within 3 days by co-culturing a glioblastoma stem cell (GSC) neurosphere (NS) with human primary astrocytes (HA), human brain microvascular pericytes (HBVPs), and human cerebral microvascular endothelial cells (hCMEC/D3) in the presence of 1.5% human serum (HS) or serum-free Neurobasal medium (NB). **(b)** Representative confocal images showing the expression of the canonical stem cell markers Nestin, SOX2, and OLIG2 in GSC neurosphere after 72-h culture under HS or NB conditions. Scale bar, 100 μm. **(c)** Quantification of Nestin-, SOX2-, and OLIG2-positive GSCs in HS or NB culture conditions from flow cytometry analysis in Extended Data Fig. 1d. The data are presented as the mean ± s.d. two-way ANOVA with Sidak’s multiple comparison test, from n=3 biological replicates. **(d)** Representative confocal z-stack reconstructions depicting the cellular organization of the BBB and BTB assembloids from 15 to 45 μm below the assembloid surface. In the BBB assembloids, the brain endothelial cells (red) encase the outer surface with associated HBVP (green), with an astrocytic (blue) core. The BTB assembloids display a notable structural rearrangement, such as reorganization of HA cells in proximity to the GSCs (white). Scale bar, 100 μm. **(e)** Quantification of GSCs in direct contact with astrocytes in BTB assembloids. **(f)** Minimum GSC-EC distances across GSC lines were quantified from 3D reconstructions and categorized into distance bins (<20 μm, <40 μm, and <60 μm). The bars represent the mean fraction of GSCs within each distance threshold. **(g)** Representative TEM images showing ultrastructure of BBB and BTB assembloids. Left panel displaying the cellular organization in the BBB and BTB assembloids. The cell types are distinguished using different colors. Scale bar, 5 μm. The magnified insets (right panels) show the formation of intact tight junctions (blue arrows) between adjacent brain endothelial cells in the BBB assembloids, while the tight junctions in the BTB assembloids appear disrupted or truncated. Scale bar, 1 μm.

Immunofluorescence (IF) staining and flow cytometry showed GSCs maintained stable SOX2, Nestin, and OLIG2 expression after 72 h in either NB or 1.5% HS, confirming preservation of their stem-like state under HS **(Fig. 1b, 1c, and Extended Data Fig. 1c and 1d)**. All GSCs in both BTB assembloids and corresponding NS showed increased proliferation rate in HS condition compared to NB (**Extended Data Fig. 1e and 1f)**.

To map cellular organization within the BTB assembloids, we differentially labeled each population: HBVPs (CellTracker Green), HAs (CellTracker Violet), hCMECs (RFP), and GSCs (Nestin staining). In BBB assembloids (no tumor), hCMECs associated with HBVPs to form a continuous outer shell expressing tight junction and BBB markers, while HAs populated the core^35^. In BTB assembloids, invasive GSCs reorganized all compartments and disrupted the endothelial surface lining (**Fig. 1d**). Measurement of astrocyte-tumor proximity showed that a substantial fraction of GSCs were in direct contact with astrocytes (85.6% GBM-X6, 78.7% GS184, 82.2% GS401) (**Fig. 1d-e**), indicating consistent astrocyte rearrangement and close physical interaction with tumor cells. This mirrors reactive astrocytosis observed in GBM tissues, which has been implicated in promoting invasion and migration^39^, suppressing immune responses^40,41^, and enhancing resistance to chemoradiotherapy^42,43^. Similarly, analysis of the spatial relationship between GSCs and ECs revealed a pronounced tumor-vascular association within BTB assembloids. Quantification of the ‘shortest distances’ demonstrated that the majority of GSCs localized very closely to the endothelium across all GSC lines, with ∼60-70% of GSCs positioned within 20 μm of ECs, increasing to >90% within 40 μm and approaching complete association (< 60 μm) (**Fig. 1f**). This perivascular enrichment indicates preferential GSC localization to the endothelial niche and supports the establishment of a vascular stem cell niche within the assembloid system. Together, these quantitative analyses validate reproducible spatial reorganization of both astrocytic and endothelial compartments around invading GSCs in the BTB assembloid and demonstrate conservation of these proximity patterns across multiple GBM patient lines.

We used transmission electron microscopy (TEM) to further examine the ultrastructure of BTB assembloids. NVU cell types were identified by established TEM criteria^35^ and as demonstrated in **Extended Data Fig. 1g**. The GSCs showed hallmark features, including polymorphic, convoluted nuclei (**Extended Data Fig. 1h-1**, blue), mitochondrial cristae breakdown (**Extended Data Fig. 1h-2**, green arrow), prominent Golgi (**Extended Data Fig. 1h-3**, red arrow), and abundant intermediate filaments and vesicles (**Extended Data Fig. 1h-3**). Consistent with our IF data, BTB assembloids (relative to BBB controls) exhibited disrupted tissue organization and a marked reduction/fragmentation of endothelial tight junctions at the surface (**Fig. 1g**).

### Permeability in BTB assembloids

GBM lesions undergo angiogenesis and vascular co-option as they grow, producing tortuous, dilated vessels and a leakier BTB than the healthy BBB^44^. To quantify tumor-induced leak in BTB assembloids, we measured dextran influx. The initial cell number for generating each GSC aggregate was tailored to their growth kinetics as neurospheres. As shown previously, BBB assembloids are impermeable to dextran^30,35^, whereas BTB assembloids display GSC density-dependent leakiness, often higher near the tumor mass, consistent with local barrier disruption (**Fig. 2a; Extended Data Fig. 2a; Supplementary Fig. 1a**). Leakiness was further elevated in 1.5% HS vs NB, likely reflecting faster GSC expansion (**Fig. 2a; Supplementary Fig. 1c**).

**Fig. 2.**
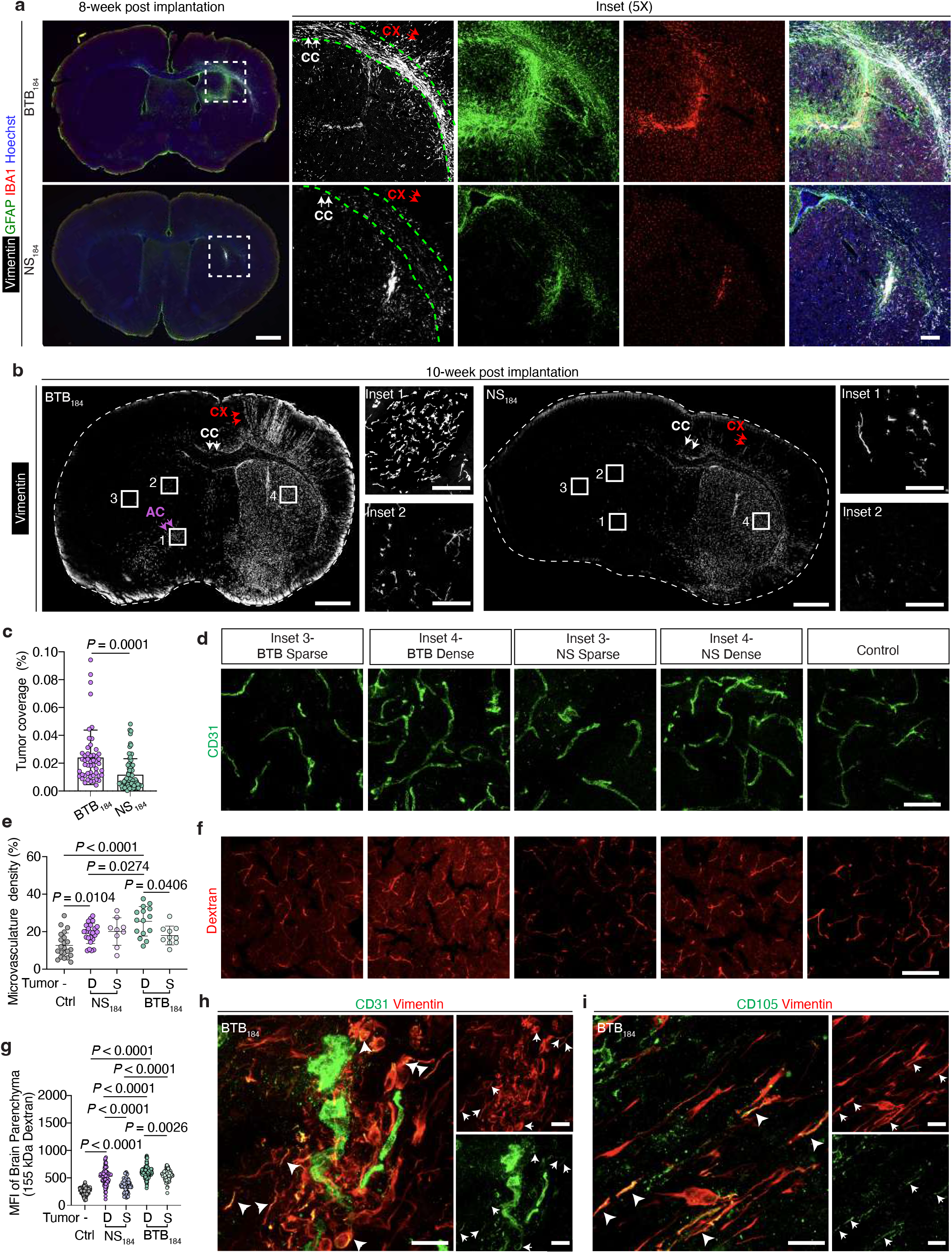
Dextran permeability and disruption of barrier integrity in BTB assembloids. **(a)** Bar graphs showing the influx of TRITC-dextran (4.4 kDa, 10 μg/mL) into the BTB assembloids in a GSC density-dependent manner in HS conditions. The data are represented as mean fluorescence ± s.d from n=3 biological replicates and n_assembloids_=3-8. The *P* values were calculated using two-way ANOVA with Tukey’s multiple comparison test. **(b)** Representative confocal images showing the expression of junction marker, ZO-1 (magenta) from immunofluorescence staining of the BBB and BTB assembloids with varying GSC (green) count. Scale bar, 100 μm. **(c)** Mean fluorescence intensity of ZO-1 expression on the brain endothelial cells in the BBB and BTB assembloids (from Fig. 2b). The data are represented as mean fluorescence ± s.d from n=3 biological replicates, n_assembloids_ = 3-8 assembloids. The *P* values were calculated using two-way ANOVA with Tukey’s multiple comparison test. Representative confocal images showing the co-staining of **(d)** VE-cadherin (white) and Glut1 (yellow), and the expression of **(e)** Mfsd2a (magenta) in the BTB and BBB assembloids. Scale bar, 100 μm. **(f, g, h)** Quantification of mean fluorescence intensity from the images in Fig. 2d and 2e, respectively. The data are represented as mean ± s.d. from n=3 biological replicates and n_assembloids_ = 4-8 assembloids. The *P* values were calculated using one-way ANOVA with Dunnett’s multiple comparisons test. **(i)** Immunofluorescence staining of COL IV in BTB assembloids and NS. Scale bar, 100 μm. **(j)** Quantitative analysis showing the GSCs expression of COL IV in assembloids *vs.* NS. The data are represented as mean ± s.d. from n_assembloids_ = 3-7 assembloids. The *P* values were calculated using one-way ANOVA with Sidak’s multiple comparisons test. **(k)** Gene expression (average of triplicate) of brain ECs isolated from BBB (EC_BBB_), GS184-BTB (EC_184_), and GS401-BTB (EC_401_) assembloids. The selected genes are related to junctions, transcytosis, transporters, angiogenesis, ECM/BM, inflammation and EndoMT. n=3 biological replicates, n_assembloids_ =1000.

To assess size selective permeability, we compared small (4.4 kDa) vs large (155 kDa) fluorescent dextrans. Entry of 4.4 kDa dextran in BTB assembloids was significantly higher than in BBB controls, whereas 155 kDa dextran remained low and near background in both groups (**Extended Data Fig. 2b**). As an orthogonal readout, we also measured the permeability of temozolomide (TMZ; a low-molecular weight and BBB-penetrant chemotherapeutic) using liquid chromatography-mass spectrometry (LC-MS) and showed detectable TMZ accumulation in both BBB and BTB assembloids (no significant difference between the two groups) (**Supplementary Fig. 1d**). Taken together, these data indicate that, despite tumor-induced leakiness, the barrier retains molecular sieving (small solutes are more permeable than large macromolecules), supporting the physiological relevance of the model for drug-delivery studies where compound size and physicochemical properties govern access.

To study the importance of direct cell-cell contact in modulating barrier disruption, we exposed BBB assembloids to GSC-conditioned medium (generated from ∼20,000 GSCs/mL; representing a cell density more than 40 times higher than that utilized in the BTB assembloids). Dextran entry did not increase (or did only modestly) even after 72 h (**Extended Data Fig. 2c; Supplementary Fig. 1b**). In contrast, BTB assembloids containing <1,000 GSCs/assembloid showed >2-fold increased dextran permeability (e.g., GS401) (**Fig. 2a**), highlighting that physical proximity, not just soluble factors, contributes to the leakier barrier.

The physical proximity of GSC and EC has been reported to create a localized perivascular niche where their interactions regulate GSC stemness, migration, therapy resistance and tumor dynamics^45^. Live confocal imaging supports this perivascular-niche mechanism. ECs are seen to elongate and migrate toward tumor clusters, sometimes extending into them (**Extended Data Fig. 2d, Supplementary video 1-4, 6**), a behavior consistent with angiogenic sprouting and EndoMT-like transitions^46,47^. In a late-stage, GSC-overgrown assembloid embedded in collagen, we observed a sequential escape: endothelial cells migrated out first, followed by GSCs, which then established microsatellite, perivascular-like niches around the primary assembloid (**Supplementary video 2**). Altogether, endothelial cells actively engage the tumor, and this direct interaction correlates with local tight-junction disruption and increased permeability.

### Disruption of barrier integrity by GSCs

We observed a GSC density-dependent loss of ZO-1 at the BTB assembloid surface, indicating tight-junction disruption (**Fig. 2b, 2c).** VE-cadherin (VE-cad), which is essential for adherens junctions and vascular permeability^48^, was likewise reduced in BTB vs BBB assembloids (**Fig. 2d, 2f**). Consistently, western blot analysis showed decreased ZO-1, VE-cad and Claudin-5 across BTB assembloids relative to BBB controls (**Extended Data Fig. 2e, 2f).** Together, these data point to tumor-driven junctional remodeling that compromises barrier integrity. We also assessed GLUT1, the principal endothelial glucose transporter and BBB marker^49^. Co-staining with VE-cad revealed lower GLUT1 on BTB endothelium than BBB (**Fig. 2d,2g**), consistent with reports of reduced GLUT1 in tumor-associated microvessels vs normal BBB^50,51^.

The lipid transporter MFSD2A (major facilitator super family domain containing 2a) is known to suppress caveolae-mediated transcytosis and maintains BBB integrity^52^. We found a marked decrease in MFSD2A expression in the BTB assembloids compared with the BBB control (**Fig. 2e, 2h**), in line with *in vivo* observations that loss of MFSD2A in tumor endothelium enhances permeability^53^.

To probe extracellular matrix (ECM) remodeling, collagen IV (COL IV) staining showed increased deposition around GSCs in the BTB assembloids vs matched NS (**Fig. 2i, 2j**), consistent with tumor-stromal interactions that reinforce basement membrane and facilitate adhesion, migration and vascular co-option in GBM^54,55^. This observation also aligns well with *in vivo* findings where GBM cells remodel the perivascular niche through excessive collagen IV accumulation to support invasion and neovascularization ^54^. Spatial fluorescence intensity profiling across the assembloids further demonstrated that COL IV enrichment localized preferentially to GSC-rich regions within the BTB assembloids, whereas the corresponding NS controls exhibited a comparatively more uniform signal distribution without focal enrichment (**Supplementary Fig. 1h**). This spatial association was consistently observed across all GSC lines examined, supporting the presence of localized ECM remodeling within the tumor-associated neurovascular niche.

Despite increased BTB “leakiness,” drug delivery can remain limited by efflux transporters, a major contributor to multidrug resistance^56^. P-glycoprotein (P-gp), which is highly expressed at the BBB and linked to chemoresistance^35,57^, is robustly expressed in BTB assembloids (**Extended Data Fig. 2g; Supplementary Fig. 1e**). Functionally, Rhodamine-123 (Rh123; a P-gp substrate) uptake assays showed higher baseline Rh123 in BTB vs BBB assembloids (reflecting greater leak), and verapamil (inhibitor for P-gp) pre-treatment significantly increased Rh123 in both models (**Extended Data Fig. 2h**). The verapamil-induced fold-increase was comparable between the BBB and BTB assembloids (especially GBM-X6 and GS184; **Supplementary Fig. 1f**), aligning with their similar P-gp levels. Thus, even in the leakier BTB assembloid, active P-gp efflux persists and can limit intracellular drug accumulation. This effect may contribute to chemoresistance in the tumor microenvironment despite compromised junctions.

### Transcriptomic profiling of brain EC in BTB assembloids

Transcriptomic profiling of the endothelial population isolated from BBB assembloids (EC_BBB_), GS184-BTB and GS401-BTB assembloids (EC_184_ and EC_401_) was performed by bulk RNA sequencing. First, benchmarking EC_BBB_ against a recently published compendium of 109 datasets across 22 libraries (including induced pluripotent stem cells (iPSC)-derived brain EC, primary endothelial (adult, fetal and human pluripotent stem cell (hPSC)-derived EC) and epithelial cells (Epi), and choroid plexus (CP) organoids)^58^, confirmed that EC_BBB_ retained a distinct endothelial identity with minimal epithelial characteristics (**Supplementary Fig. 1g**), supporting the BBB fidelity of our assembloids.

In EC_BTB_, we observed a downregulation in several tight junction/adherens junction-associated genes (such as *TJP1*, *CLDN5*, *JAM2*, *CDH5*, *PECAM1, ESAM*) and *MFSD2A* compared to the control EC_BBB_ (**Fig. 2k**), consistent with our IF staining showing decreased Mfsd2a protein expression (**Fig. 2h**). Mfsd2a functions to suppress caveolae-mediated transcytosis, and loss of Mfsd2a leads to an upregulation of caveolae vesicle trafficking^52^. Concordantly, in EC_BTB_, integral membrane proteins caveolin-1 (*Cav1)* and caveolin-2 (*Cav2)* were upregulated (**Fig. 2k**). Transport programs were also remodeled – several SLC transporters linked to GBM metabolism (*SLC43A3*^59^*, SLC16A1*^60^*, SLC7A5*^61^) and efflux pumps (*ABCA1*)^62^ were elevated (**Fig. 2k**), in line with reports that endothelial transport reprogramming supports GBM progression and chemoresistance^63,64^.

Markers of endothelial activation/neuroinflammation in the EC_BTB_ increased, including *ICAM1* and *VCAM1* (mediate leukocyte recruitment) together with cytokines *IL-1*, *TNF-α, IL-6*, and *CCL2* (**Fig. 2k**), a profile associated with enhanced BBB permeability^65^. We also detected endothelial-to-mesenchymal transition (EndoMT)-like shift, which plays an important role in chemoresistance and promoting tumor progression^66^. Upregulation of mesenchymal markers such as *IL-1β*, collagen1α2 (*COL1A2*), SLUG (*SNAI2*), and *TGFβ2*, with concomitant downregulation of endothelial marker such as CD31 (*PECAM1*) and VE-cadherin (*CDH5*) (**Fig. 2k**) is consistent with EndoMT signatures in GBM^46,66,67^. Finally, ECM-remodeling genes (such as collagens, agrin *(AGRN)*, tenascin-C (*TNC)*, fibronectin (*FN1*), and nidogen-1 (*NID1*)) were increased (**Fig. 2k**). This aligns with increased collagen IV level around the tumor regions (**Fig. 2i, 2j**) to support tumor invasion and angiogenesis^54,55^. Altogether, these endothelial changes in the BTB context indicate that the brain vasculature not only loses barrier integrity but actively shifts from a ‘barrier-maintaining’ to a ‘pro-tumor’ state (loosening junctions, enhancing transcytosis/transport, activating inflammatory programs, and remodeling matrix).

### GSC_BTB_ display increased invasiveness and mesenchymal gene signature

To profile tumor-endothelial interactions, bulk RNA-sequencing was also performed on GS184, GS401 and EC isolated from the BTB assembloids. More than 1000 assembloids were pooled into a single sample to ensure RNA yield (**Fig. 3a)**, with n=3 biological replicates. First, GSCs and ECs were separated from astrocytes/pericytes by fluorescence-activated cell sorting (FACS) based on low PDGFRβ expression (GSC and EC), and then distinguished by high-EGFR (GSC) vs low-EGFR (EC) (**Extended Data Fig. 3a**). The EC population isolated from the BBB assembloids (EC_BBB_), and GSC neurospheres (NS_184_ and NS_401_) were used as controls. PCA showed clear separation between BTB-derived GSCs (BTB_GSC_) and their neurosphere counterparts (NS_GSC_), and between EC_BTB_ and EC_BBB_ (**Extended Data Fig. 3b**), indicating strong coculture-dependent reprogramming. Differential gene-expression analysis identified 1,167 upregulated and 661 downregulated genes in GSCs from BTB_184_ relative to NS_184_ (**Fig. 3b**), while GSCs from BTB_401_ displayed 590 upregulated and 44 downregulated genes relative to NS_401_ (**Fig. 3c**).

**Fig. 3.**
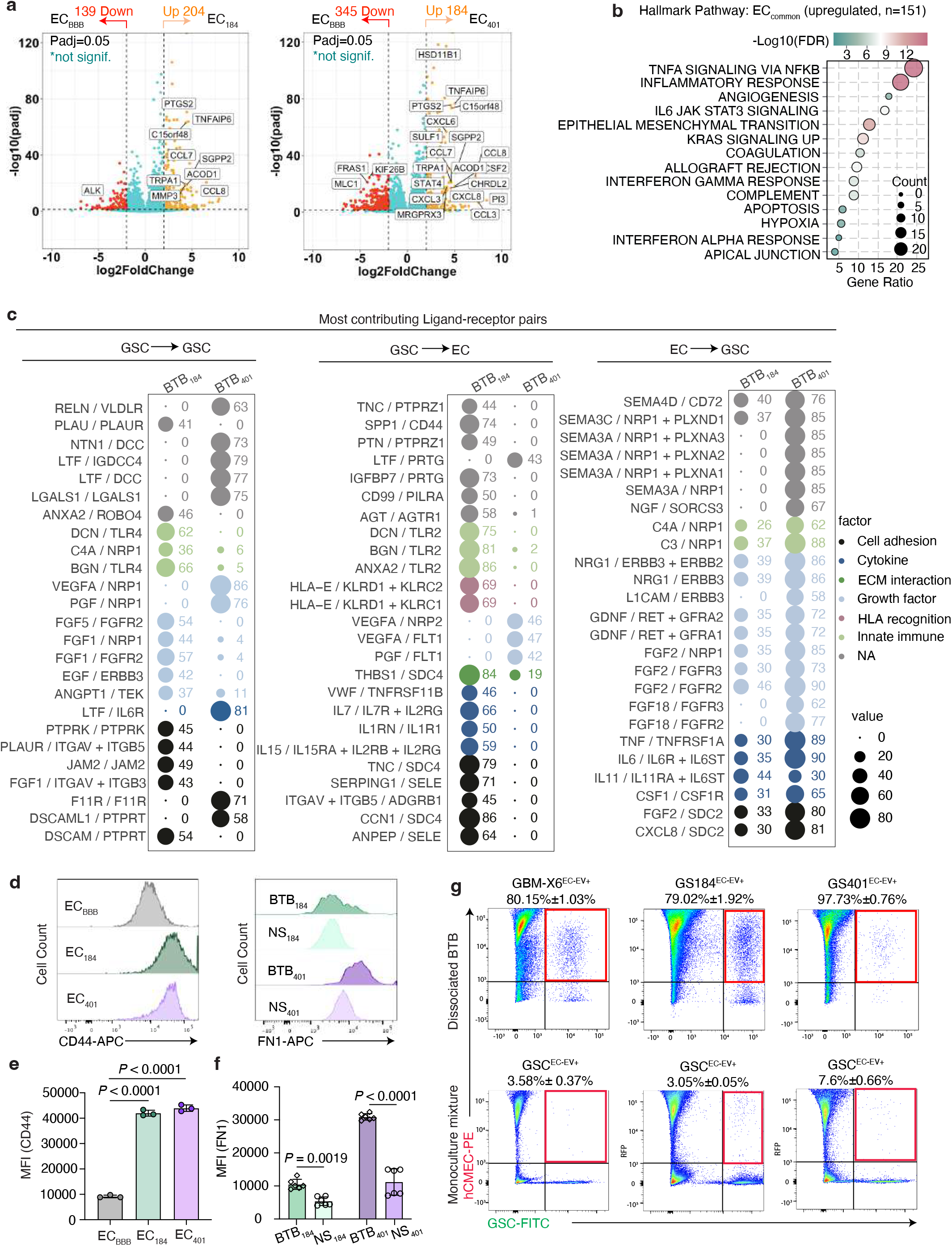
Transcriptomic profiling of GSCs shows enhanced tumor aggressiveness in BTB assembloids. **(a)** Schematic of fluorescence-activated cell sorting (FACS) workflow. **(b, c)** Volcano plot showing differential expression of genes between **(b)** BTB_184_ vs NS_184_; **(c)** BTB_401_ vs NS_401_. **(d)** Enriched KEGG pathways in BTB_184_ and BTB_401_. (False discovery rate (FDR) < 0.05). Commonly enriched pathways in both BTB_184_ and BTB_401_ are highlighted in red. **(e)** Gene ontology chord plot showing the top 10 enriched biological processes (BP) for GSC_common_ and the genes involved in each BP. **(f)** Bubble heatmap displaying gene expression fold change (FC) in the GSCs of BTB_184_ and BTB_401_ assembloids compared to the respective controls, NS_184_ and NS_401._ Genes associated with stemness, epithelial-mesenchymal transition (EMT), invasion/migration, angiogenesis, hypoxia, inflammatory response, and ECM remodeling were compiled from the literature^95,96^. **(g)** Measurement of cell invasion using the transwell invasion assay (refer to **Extended Data** Fig. 3j) comparing between the BTB_GSC_ and NS_GSC_ groups. GSC invasion is tracked using time-lapse imaging. Quantification of the total number of GSCs that have invaded to the basal side of the transwell insert shows that BTB_GSC_ are more invasive than NS_GSC_. The data are represented as mean ± s.d. from n=3 biological replicates, n_assembloids_ = 8. The *P* values were calculated using one-way ANOVA with Dunnett’s multiple comparisons test. **(h)** Flow cytometry analyses of protein expression of the representative genes related to stemness, EMT, invasion and inflammation (SOX2, Nestin, PDGFRA, MET, CD44) in GSC_BTB_ and GSC_sphere_. **(i)** Quantification of mean fluorescence **(from h)** demonstrates enhanced mesenchymal and self-renewal features in BTB_GSC_ compared to NS_GSC_. The data are represented as mean ± s.d. from n=3 biological replicates, n_assembloids_ = 60 dissociated assembloids. The *P* values were calculated using two-way ANOVA and Sidak’s multiple comparison test, \*\**P* < 0.01, \*\*\*\**P* < 0.0001).

Pathway analysis revealed shared upregulation of cytokine-cytokine receptor, ECM-receptor, protein digestion/absorption, focal adhesion, PI3K-AKT and MAPK signaling pathways (**Fig. 3d**), which are programs linked to cytoskeletal reorganization and cell locomotion, critical for promoting GBM cell adhesion, proliferation, motility and apoptosis resistance^68,69^. Line-specific differences were also evident: GSCs from BTB_184_ (primary) showed enrichment in inflammatory-related pathways such as TNF, NF-κB, and IL-17 signaling pathways known to promote tumor invasion^70^, while GSCs from BTB_401_ favored EGFR tyrosine kinase inhibitor resistance, calcium signaling pathway and Ras signaling pathway that are predominantly involved in tumor progression, angiogenesis and resistance to drugs^71,72^.

We identified 143 genes upregulated in both BTB lines (GSC_common_) compared to their respective NS_GSC_ (**Extended Data Fig. 3c)**. Gene Ontology (GO) analysis for these common genes showed several enriched biological processes, including those related to ECM organization, extracellular structure organization, angiogenesis, and blood vessel morphogenesis (**Fig. 3e**), with epithelial-mesenchymal transition (EMT) identified as the most enriched hallmark (**Extended Data Fig. 3d**). Correspondingly, TNF-α/NF-κB signaling, which are key EMT drivers^73,74^ was also highly elevated. Downregulated programs differed by line: BTB_184_ showed loss of neurodevelopment, neurogenesis and cell differentiation, suggesting that exposure to vascular and glial cues induces a dedifferentiated, stem-like state characterized by reduced axonal growth and neuronal specification (**Extended Data Fig. 3f**). In contrast, BTB_401_ downregulated tissue remodeling, negative regulation of angiogenesis, and vascular morphogenesis, suggesting a shift toward a pro-angiogenic, invasive phenotype, consistent with the mesenchymal features of this tumor subtype. In line with prior GSC taxonomy, where constructive GSCs are enriched for developmental pathways related to neuron, brain and glial cells, while invasive GSCs are driven by enrichment of ECM and angiogenesis pathways^75^, these data indicate that NVU cues reprogram GSCs toward the invasive state over the constructive state.

To further dissect subtype-specific adaptations, we compared BTB_401_ with BTB_184._ This comparison showed enrichment of hallmark pathway, including E2F targets^76,77^, G2M checkpoint^78,79^, MYC targets^80^, DNA repair and oxidative phosphorylation in BTB₄₀₁ (**Extended Data Fig. 3g**), indicating enhanced proliferative and metabolic activity of a more aggressive and cell cycle-driven phenotype. This is confirmed by Kyoto Encyclopedia of Genes and Genomes (KEGG) pathway analysis (**Extended Data Fig. 3e**) showing enhanced cell cycle, DNA replication and oxidative/glycolytic metabolism. Correspondingly, BTB₄₀₁ upregulated core cell-cycle regulators (*E2F1/2, CDC20, CDK1, CCNB1* and *TTK*) and DNA-repair components (*RAD51, POLA2, LIG1* and *RFC5*) (**Extended Data Fig. 3h, 3i**). Together, these findings indicate that BTB₄₀₁ exhibits a highly proliferative, genome-maintenance, metabolically flexible program that is consistent with aggressive, treatment-resistant and mesenchymal-dominant GBM.

### GSC_BTB_ exhibit enhanced features seen in clinical GBM samples

We next examined key regulatory genes curated from two public GBM patient datasets (GSE145645 and GSE147352) from Gene Expression Omnibus (GEO). Differential expression analysis confirmed that these genes are enriched in GBM relative to low-grade glioma (LGG) and normal brain tissue (**Supplementary Fig. 2a and 2b**). Consistently, several of these GBM-prevalent genes were upregulated in BTB_GSC_ compared to NS_GSC_ (**Fig. 3f)**, spanning programs related to stemness, EMT, invasion/migration, angiogenesis, hypoxia, inflammatory response and ECM remodeling^81–83^.

Hallmark genes related to stemness, EMT and invasion were upregulated in BTB_GSC_, indicating that NVU co-culture drives a mesenchymal-associated, more invasive and self-renewing state. We also observed ECM remodeling in BTB_GSC_ with elevated *FN1, TNC, CD44 and COL4A1*) (**Fig. 3f**), in line with GBM’s aberrant ECM rich in fibronectin, tenascin-C, hyaluronan^84,85^ and fibrillar collagen^86^. These changes are consistent with ECM-supported tumor growth, invasion and angiogenesis^55^. Functionally, we used transwell (Matrigel) invasion assays to show significantly more BTB_GSC_ (than NS_GSC_) traversed through the Matrigel into the basolateral chamber (using human serum as a chemoattractant) (**Fig. 3g; Extended Data Fig. 3j**). Flow cytometry further revealed higher expression of several markers in the BTB_GSC_ (vs NS_GSC_) that are known to promote GBM aggressiveness, including *SOX2*^87,88^*, NESTIN*^88,89^, *PDGFRA*^90,91^, *MET*^92,93^ and *CD44*^94^ (**Fig. 3h and 3i)**.

### Transcriptional reprogramming of EC_BTB_ bolsters tumor progression

The dynamic GBM-microvascular interaction disrupts tissue homeostasis and promotes tumor progression^97^. DEG analysis of the EC revealed 204 and 184 upregulated genes in BTB assembloids established with GS184 (EC_184_) and GS401 (EC_401_), respectively compared to control EC_BBB_ (**Fig. 4a**). While only ∼8% of upregulated genes in GSC_common_ overlapped (**Fig. 3e**) signifying notable inter-heterogeneity, ∼64% of the upregulated genes overlapped between EC_184_ and EC_401_ (EC_common_) (**Extended Data Fig. 4a)**, indicating a homogeneous endothelial response to the GSCs.

**Fig. 4.**
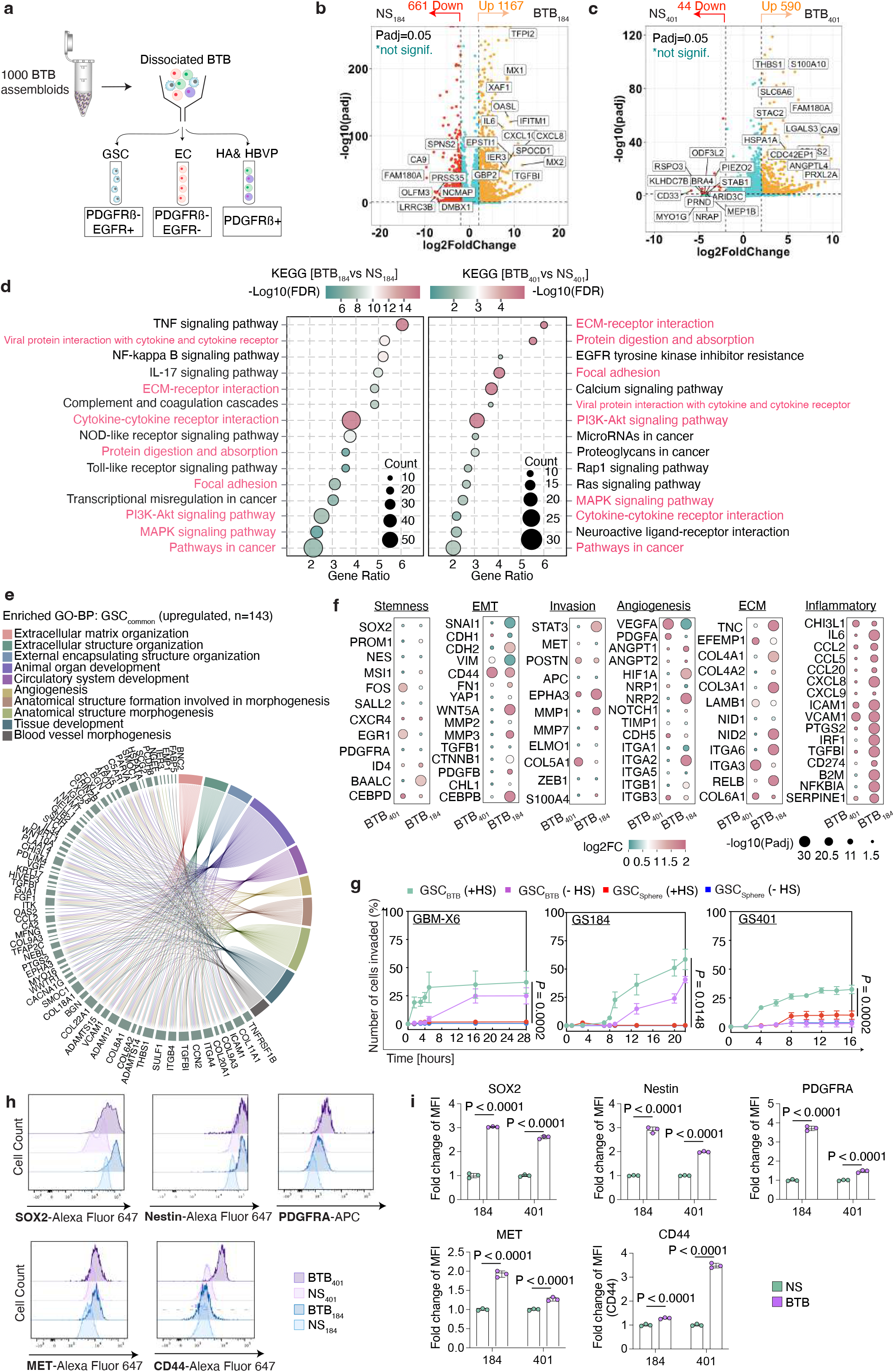
Transcriptomic analysis showing genetic alteration of the endothelial cell population in BTB assembloids. **(a)** Volcano plot showing differential gene expression for EC _BTB-184_ and EC _BTB-401_. **(b)** Enrichment analysis of the upregulated EC_common_ showing the top enriched hallmark pathways (FDR < 0.05). **(c)** Inferred ligand-receptor interactions through applying the ICELLNET framework^105^ to the bulk transcriptomic data of GSC_BTB_ and EC_BTB_. The plots display the top 25 GSC autocrine interactions (left), as well as GSC_ligand_-to-EC_receptor_ and EC_ligand_-to-GSC_receptor_ paracrine communication, along with their individual interaction scores. Only regulated genes (Log2FC >1) from each cell type were extracted and applied. Other genes were input with value 0. Genes associated with ligands and receptors in the database are identified and normalized by the maximum gene expression observed for each gene. These normalized values are then used to calculate a communication score between the two cell types. **(d)** Flow cytometry analysis of EC and GSC in GS184- and GS401-BTB assembloids. *Left:* CD44 surface expression on EC from BBB controls and BTB. *Right:* FN1 protein levels in GSCs isolated from GS184-BTB and GS401-BTB assembloids. **(e)** Quantification of CD44 median fluorescence intensity in EC_BBB_, EC_184_ and EC_401_. **(f)** Quantification of FN1 median fluorescence intensity in GSCs from BTB assembloids compared to NS. **(g)** Flow cytometry analyses demonstrating a high proportion of GSCs (80% of GBM-X6; 79% of GS184; 98% of GS401) have positive RFP signal (GSC^EC-EV+)^ in the BTB assembloids. Cell gating is adjusted based on the control group composed of the RFP-expressing hCMEC/D3, HBVP, HA and GSC monocultures mixed together in the same ratio as the assembloid co-culture group immediately before analysis. The experiment was conducted in triplicates and each BTB sample contains 60 dissociated assembloids.

Pathway analyses of EC_common_ (upregulated genes) in BTB assembloids identified inflammation-related pathways including IL-17, TNF and NF-κB signaling pathways^98^ (**Extended Data Fig. 4b**), consistent with BBB disruption and injury response. Hallmark pathway analysis also revealed enrichment of EMT and angiogenesis pathways in EC_common_ (**Fig. 4b**). Indeed, inflammatory GBM microenvironment can drive EMT in endothelial subsets^99^, contributing to resistance to anti-angiogenic therapy^100^, chemoresistance^67^ and radio-resistance^101^. Hypoxia pathways were also enriched in EC_common_, aligning with pro-angiogenesis activation (**Fig. 4b**). Conversely, GO analyses revealed several overlapped downregulated pathways between EC_184_ and EC_401_ (EC_down_) relative to control EC_BBB_, including axon development, neurogenesis, nervous system development and neuron differentiation (**Extended Data Fig. 4c**). This is consistent with reports that BBB perturbation leads to neuronal dysfunction, neuroinflammation and neurodenegeneration^97,102^. Finally, when benchmarked against a set of 78 genes upregulated in malignant GBM vasculature in patients^103^, 46 (in EC_184_) and 40 (in EC_401_) genes were upregulated compared to control EC_BBB_ (**Extended Data Fig. 4d**). This shows stronger concordance than a recent study^51^ that identified only 24 (out of the 78) upregulated genes in ECs from primary and secondary human GBM compared to control vessels.

### GSC-EC crosstalk and ligand-receptor networks

Reciprocal signaling between GSC and EC is crucial for regulating GSC stemness, migration, invasion, angiogenesis and therapeutic resistance^69,104^, and is mediated by a complex interplay of soluble factors, including growth factors, cytokines, chemokines and EVs^48^. Indeed, GO analysis of EC_common_ genes showed enrichment of cytokine/chemokine activity and receptor-ligand interaction pathways compared to the control EC_BBB_ (**Extended Data Fig. 4e**).

To map these communications, we reconstructed both autocrine and paracrine ligand-receptor (L-R) networks between GSCs and ECs. Using ICELLNET^105^, we quantified potential L-R scores based on differentially expressed upregulated genes, which showed growth factors and cell adhesion molecules as dominant mediators (**Extended Data Fig. 4f**, **4g)**. The top 25 L-R pairs revealed extensive growth factor-driven signaling and cell-ECM interactions (**Fig. 4c**). Several GBM autocrine interactions were also evident, including *VEGFA - NRP1*^106,107^, *EGF - ERBB3*^108^ and *FGF1/5 - FGFR2/3*^69^, all of which have been previously implicated in maintaining GSC stemness, promoting vascular co-option, and enhancing therapy resistance (**Fig. 4c**).

In addition to these self-sustaining loops, the GSC→EC interactions were enriched for pro-angiogenic and ECM-remodeling signals such as *TNC-PTPRZ1, SPP1-CD44*^109,110^*, BGN/DCN-TLR2 and VEGFA-NRP2*^69^ (**Fig. 4c**), indicating that GSCs actively impose a pro-inflammatory, adhesive and angiogenic state on the endothelial cells, thereby shaping a tumor-permissive vascular niche. Conversely, the EC→GSC communication were characterized by endothelial-derived trophic and cytokine factors, including FGF2-FGFR2/FGFR3^111^, IL6-IL6R/IL6ST^112^ and CXCL8-SDC2^113^, which have been implicated in promoting GSC proliferation, EMT and BTB remodeling. Together, these interactions delineate a dynamic and reciprocal communication underlying the vascular adaption of GSCs in the BTB.

As complementary validation, we cross-referenced upregulated ligands/receptors against the CelltalkDB curated database of experimentally validated L-R interactions^114^ (**Supplementary Fig. 3**). This corroborated many of the ICELLNET-predicted pairs and revealed both conserved and subtype-specific communication patterns across both BTB_184_ and BTB_401_ assembloids (**Supplementary Fig. 3a-3d**). The L-R matrices highlighted shared growth factor (*FGF2-FGFR2*, *VEGFA-VEGFR*) and ECM-related (*BGN-TLR4*, *COL4A1-ITGA*) interactions, alongside subtype-enriched signaling pairs such as *IL6-IL6R* and *CXCL8-SDC2*, and several collagen-integrin interactions associated with mesenchymal transition and vascular remodeling. Collectively, these define a complex and dynamic GSC-EC communication landscape that may potentially contribute to differential vascular phenotypes and therapeutic responses in the BTB.

We examined an ECM signaling pair central to the neurovascular niche and identified FN1 (on GSCs) -CD44 (on endothelial cells) as a candidate interaction for functional investigation (**Supplementary Fig. 3a, 3b**). CellTalkDB analysis highlighted upregulated FN1-CD44 in both GS184- and GS401-BTB. We then examined this at the protein level in BTB assembloids. Using the same cell-sorting gates as in our RNA analysis (**Extended Data Fig. 3a**), we measured CD44 on endothelial cells and FN1 on GSCs. In both assembloid models, endothelial CD44 was significantly higher than in BBB controls, matching the transcriptomic prediction (**Fig. 4d, 4e**). Likewise, FN1 on BTB_GSC_ was higher than in their matched neurospheres (**Fig. 4d, 4f**), consistent with the ECM remodeling programs inferred from our L-R maps. Together, protein-level analysis confirmed increased FN1 and CD44 expression in the corresponding cellular compartments, providing orthogonal support for the transcriptomically predicted FN1-CD44 candidate interaction.

### Endothelial-derived extracellular vesicles (EV) uptake by GSCs

EVs play important roles in mediating the intercellular communication between GSC and EC. In the BTB assembloids, we observed substantial internalization of EC-derived EV by GSCs using high-resolution confocal microscopy (**Extended Data Fig. 4h, Supplementary video 4, 5, 6, 7**). Flow cytometry further revealed robust transfer of endothelial RFP signal to tumor cells: 80% of GBM-X6; 79% of GS184; 98% of GS401 became RFP-positive following co-culture with RFP-expressing ECs (**Fig. 4g**). In contrast, <4% of GBM-X6 and GS184, and <8% of GS401 were RFP-positive in the control group (comprised of GSC and each NVU cell type mixed in the same proportion as the assembloids immediately before analysis). These findings are consistent with EC-to-GSC transfer of RFP-positive material at the tumor-vascular interface, spatially enabled by the close proximity of endothelial and tumor compartments within the assembloid. While the functional consequences of this transfer and the specific contribution of EVs to BTB-associated phenotypes remain to be established, these observations highlight the potential of the platform for future mechanistic investigation of EV-mediated tumor-vascular crosstalk.

### Neurovascular niche-driven reprogramming and increased BTB_GSC_ aggressiveness *in vivo*

Orthotopic GBM xenografts were established by intracranial injection of dissociated single cells from BTB_184_, NS_184_ or control BBB (NVU-only) assembloids (**Extended Data Fig. 5a**). For NS_184_, dissociated GS184 cells were mixed with each NVU cell types in the same ratio as the assembloids immediately before implantation. Tumors were established in the right frontal lobe and grown for 8-10 weeks. At 8 weeks, IF staining of human vimentin confirmed GS184 engraftment in both the BTB_184_ and NS_184_ groups (**Fig. 5a)**. To validate vimentin as a GBM marker, we also co-stained with human-specific nestin, where both markers showed strong spatial overlap across all tumor regions (**Supplementary Fig. 4a and 4b**)

**Fig. 5.**
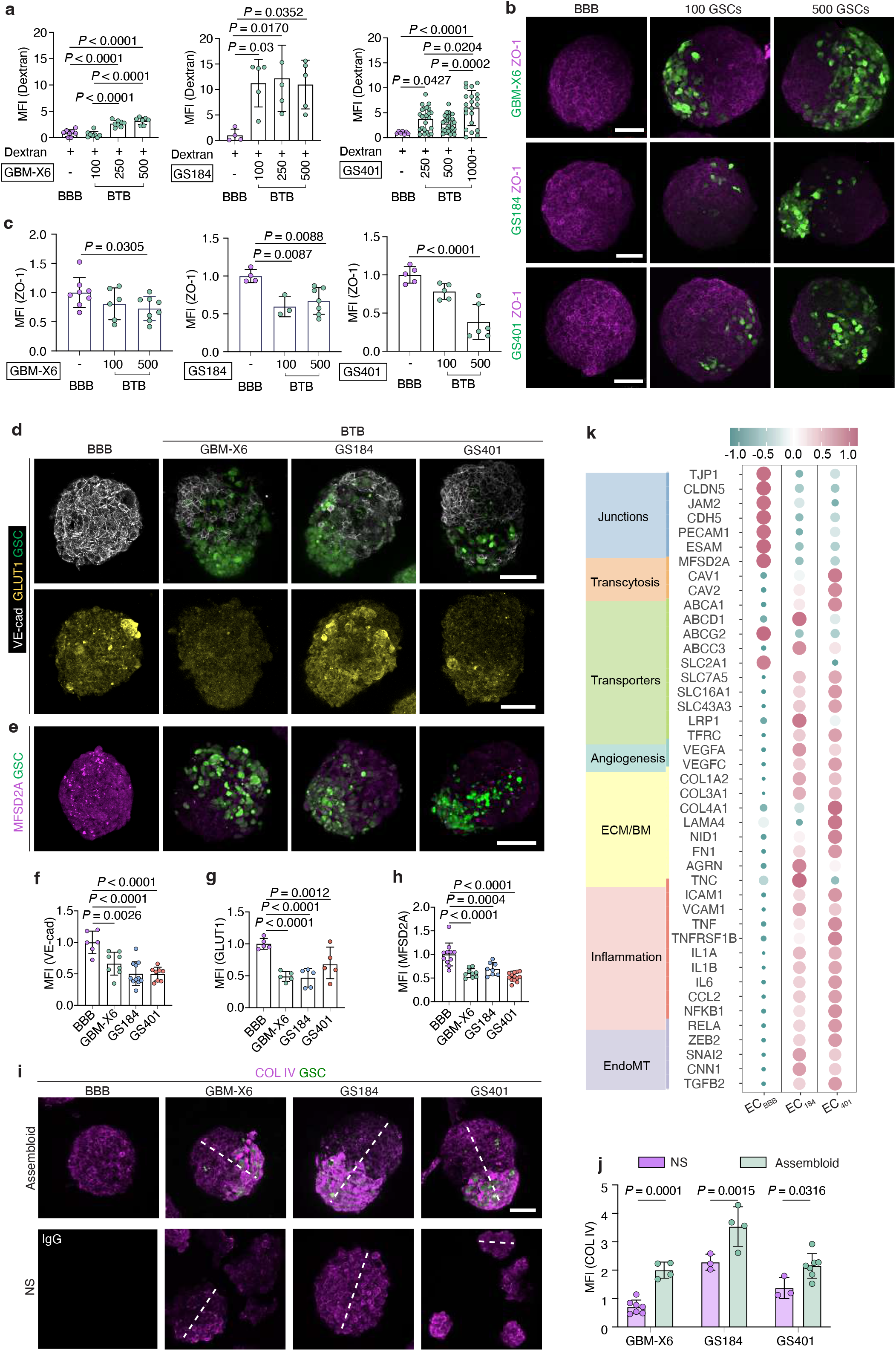
GSCs from BTB assembloids are more aggressive than the NS in orthotopic PDX mouse models. **(a)** Immunofluorescence staining of coronal brain cryosections showing engraftment of GSCs from BTB_184_ and NS_184_ in mice at 8 weeks. Whole brain sections are stained with Vimentin, GFAP and IBA1 to visualize the GSC, astrocytes, and GBM associated macrophages and microglia (GAM), respectively. Insets (5X) showing the magnified areas specified in the whole brain slices (left). CC: corpus callosum (white arrows), CX: cortex (red arrows). Green dash line outlines the CC region. Scale bar, whole brain sections: 1mm; inset: 200 μm. **(b)** Invasion of GSCs from BTB_184_ and NS_184_ at 10 weeks post-implantation in mice brain. AC: anterior commissure (purple arrows). Insets 1 and 2 show the magnified views of selected regions in the contralateral hemisphere of the brain section. GSCs from BTB_184_ exhibits higher invasion through the white matter tracts CC and AC, and into the cortex and contralateral hemisphere. Scale bar, whole brain sections: 1mm; inset: 100 μm. **(c)** Graph showing the percentage of GSC coverage in BTB_184_ and NS_184_ in mice brain. Results are quantified from 15-20 serial coronal cryosections per mouse (n = 3-4 mice each group, unpaired t test with Welch’s correction). **(d)** Representative fluorescence images and quantification **(e)** of microvasculature density (CD31 staining) in the BTB_184_ and NS_184_ groups. Images showing magnified view of insets 3 and 4 outlined from panel (B) represent tumor-sparse and tumor-dense regions in the BTB_184_ and NS_184_ groups, respectively. Scale bar, 50 μm. (n = 3-4 mice per group, the *P* values were obtained using one-way ANOVA with Tukey’s multiple comparisons test). **(f)** Representative confocal images and quantification **(g)** showing the perivascular signal of TRITC dextran in the BTB_184_ and NS_184_ groups. No significant change in dextran permeability is observed in the tumor-sparse areas compared with the control (healthy brain with no tumor). A significant increase in dextran signal outside the vessels is detected in the tumor-dense regions, with the highest leakage observed in the BTB_184_ group. Scale bars, 100 μm. (n = 3-4 mice each group, the P values were calculated using one-way ANOVA with Tukey’s multiple comparisons test). **(h, i)** Representative images showing endothelial transdifferentiation in the BTB_184_ group. Immunofluorescence staining of brain cryosections for the tumor cells with human vimentin (red) with CD31 and CD105 (green). Arrows (white) indicate colocalization of vimentin with CD31(h) and CD105 (i). Scale bars, 20 μm.

Because NVU cells can also express human vimentin (**Extended Data Fig. 5b**), as a control, we implanted dissociated cells from BBB-only assembloids in the same manner. We detected no vimentin signal throughout the brain cryosections (**Extended Data Fig. 5c**), indicating that the human NVU cells do not survive/propagate in the mouse brain, and confirming human vimentin as an effective marker for visualizing the GSC population in our assays.

By 8 weeks post-implantation, a significant level of BTB_184_-derived GSC had migrated along the corpus callosum (CC) and invaded into the cortex (CX), whereas NS_184_ tumors remained more compact with significantly fewer cells in the CC and CX (**Fig. 5a, Extended Data Fig. 5c**). By 10 weeks, BTB_184_ tumors showed greater hemispheric spread and white matter tract invasion (including the CC and anterior commissure (AC), extending to the CX and the contralateral hemisphere (**Fig. 5b**). Quantification across several cryosections showed larger tumor area in BTB_184_ vs NS_184_ (**Fig. 5c)**, indicating a more aggressive and infiltrative phenotype. IF staining revealed elevated phospho-vimentin in BTB_184_ compared to NS_184_ (**Supplementary Fig. 4c and 4d**). As phospho-vimentin marks active filament reorganization that supports cell motility and mesenchymal transition^115^, these data implicate endothelial/perivascular cues in licensing a more migratory, invasive and mesenchymal-associated state in BTB-derived GSCs.

We next assessed astrocytic activation through IF staining for GFAP^116^. Astrocytes are known to be activated by glioma cells^117,118^, characterized by hypertrophy and high GFAP expression. They directly associate with GBM cells to aid tumor progression, mesenchymal transition and therapeutic resistance^42,116,119^. Reactive, GFAP⁺ astrocytes populated peritumoral zones in both groups (**Fig. 5a and Extended Data Fig. 5c)**, with significantly higher GFAP in the BTB_184_ group (**Supplementary Fig. 4f**). Spatial analysis showed majority of astrocytes located in close proximity to GSCs in both groups (**Supplementary Fig. 4e**), consistent with earlier *in vitro* results showing astrocytes relocalize near GSC clusters in BTB assembloids (**Fig. 1d and 1e**).

Concurrent activation of astrocytes and microglia have been recognized for their pivotal role in promoting GBM malignancy and invasiveness^120^, and GBM-associated macrophages and microglia (GAMs) are more enriched in mesenchymal GBM^121,122^. Indeed, immunostaining for IBA1 showed significantly higher level of activated microglia clustering around BTB_184_ tumors compared to NS_184_ (**Fig. 5a, Extended Data Fig. 5c**), aligning with literature linking astrocyte-microglia co-activation and mesenchymal GBM to enhanced invasiveness. Together, these data indicate that BTB-derived GSCs engage the neurovascular niche to drive cytoskeletal remodeling, glial activation and invasive spread *in vivo*.

### GSCs from BTB assembloids exacerbate BBB leakage and neovascularization

We further examined vascular changes in brain regions with dense versus sparse tumor cell populations (**Fig. 5d**). IF staining for CD31 revealed that, in both the BTB_184_ and NS_184_ groups, tumor-dense regions displayed more disorganized, tortuous vasculature and microvasculature density (MVD) than contralateral tumor-sparse areas and healthy brain controls (**Fig. 5d, 5e**). To assess barrier function, we injected TRITC–dextran intravenously. BTB_184_ group exhibited greater parenchymal dextran accumulation, indicative of enhanced vessel permeability and BBB compromise compared to the NS_184_ group, especially in tumor-dense areas (**Fig. 5f, 5g).** In healthy brain tissues and tumor-sparse regions, dextran signal remained confined within the blood vessels, consistent with an intact BBB (**Fig. 5f**).

A subset of BTB_184_ tumor cells expressed the endothelial markers CD31 and CD105 (**Fig. 5h, 5i**), consistent with acquisition of an endothelial-like phenotype^123^, a process known to be important for supporting tumor growth, invasion and resistance to anti-angiogenic therapies^124^. Together, these findings show that prior conditioning by the neurovascular niche drives GSCs toward a more invasive, mesenchymal and endothelial-like state, thereby exacerbating neovascularization and BBB dysfunction *in vivo*.

### Modeling therapeutics delivery and efficacy

#### GBM targeting using BTP-7 peptide

To validate the BTB assembloids for therapeutic testing, we evaluated BTP-7, a peptide that targets the deglycosylated brevican (dg-BCAN) isoform^36^. The ECM glycoprotein Bcan presents a unique de-glycosylated isoform dg-Bcan that is selectively present in human high-grade gliomas^36,125^, and absent from healthy central nervous system (CNS) and other non-cancerous neuropathological tissues. Prior work established that BTP-7 *i)* binds dg-Bcan recombinant protein specifically, *ii)* targets GBM tumors in patient-derived xenograft (PDX) mouse models, *iii)* is specifically taken up by GSCs compared to other NVU cells, and *iv)* crosses the BBB in both human BBB assembloids and in mice^33,36^.

In our cohort, western blot analyses confirmed dg-Bcan expression across all patient-derived GSC models (**Extended Data Fig. 6a**). Following incubation and washout of Cy5-labeled BTP-7 with the BTB assembloids, we observed preferential retainment in the GSCs (**Fig. 6a**), with faster clearance from the NVU cells in both the BTB and BBB assembloids (**Fig. 6b, 6c, Extended Data Fig. 6b**). These results are consistent with our prior *in vivo* findings demonstrating effective targeted delivery of BTP-7 to orthotopic patient-derived GBM tumors^33,36^.

**Fig. 6.**
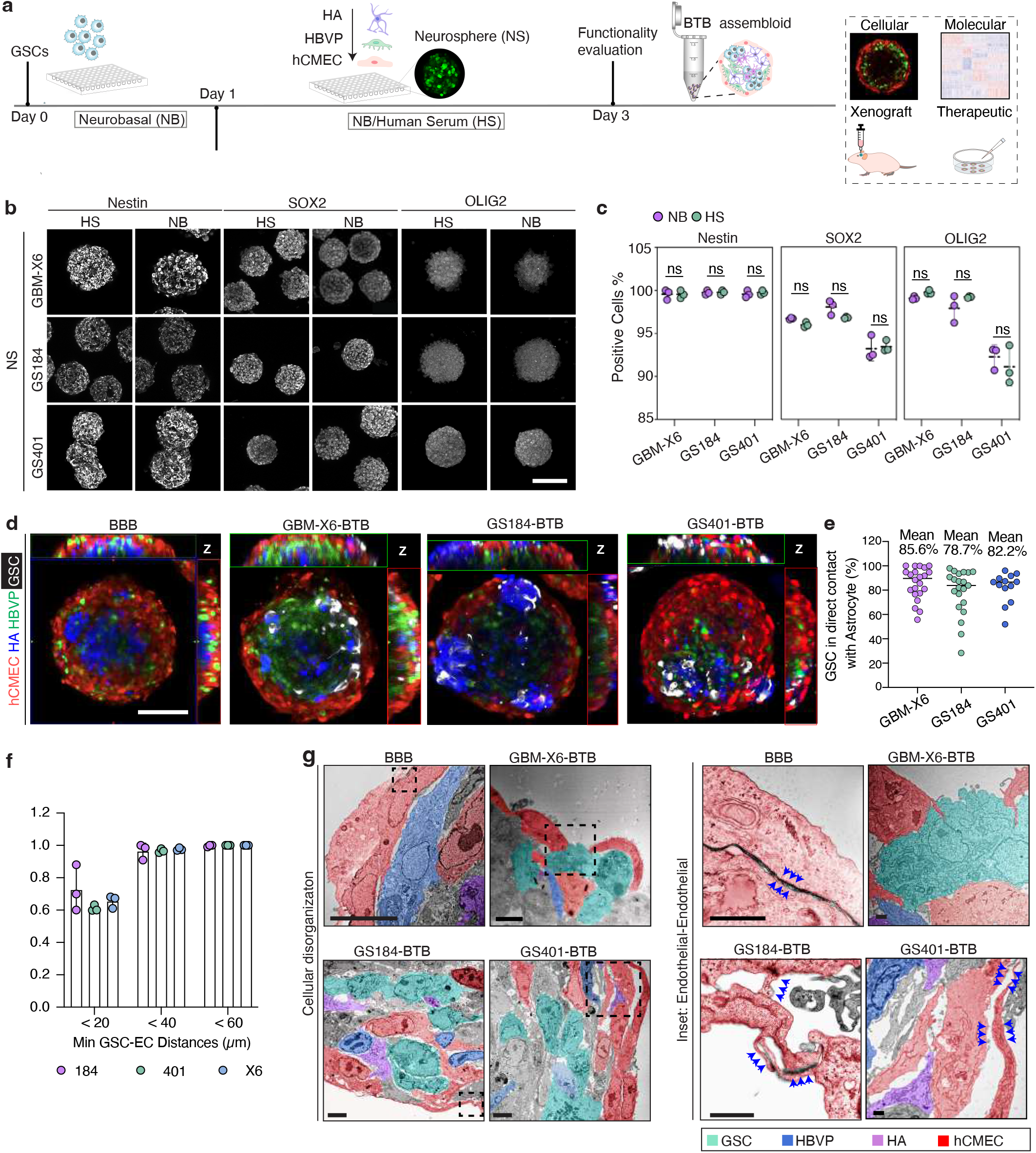
Modeling therapeutic delivery and efficacy using BTB assembloids. **(a)** Schematic showing the workflow of BTP-7 peptide targeting in BTB assembloids. Representative fluorescence images **(b)** and quantification of mean fluorescence intensity **(c)** displaying the distribution of BTP-7-Cy5 (white) in BBB and BTB assembloids over a washout period (refer also to Extended Data Fig. 6b). After 4h-incubation with BTP-7-Cy5, the assembloids are washed for a period starting at 0 h (up to 60 h). BTP-7-Cy5 is specifically retained in the GSCs over time, while displaying greater clearance rate in the NVU cells. The data are presented as mean ± s.d. from n = 3 biological replicates and n_assembloids_ = 6-8. The *P* values were calculated using two-way ANOVA with Dunnett’s multiple comparison test. Scale bar, 100 μm. **(d)** Timeline of AAV.CPP.16-GFAP-TK1 therapy combined with Ganciclovir (GCV) in BTB assembloids. Assembloids are treated with AAV (1 × 10^9^ vg/assembloid), followed by addition of 8 μM GCV after 48 h until day 9, with GCV refreshed every day. BTB assembloids were incubated with GCV for 8 h and 16 h in fresh medium. **(e)** Schematic illustration of the workflow used to quantify GBM cell growth in BTB assembloids. Following treatment, BTB assembloids were collected and dissociated into single cells using papain digestion (10-15 minutes at 37 °C). Dissociated cells were then plated for imaging and quantification of GFP⁺ GSCs, representing GBM cell populations. The total number of GFP⁺ cells was used to assess relative GBM cell proliferation or therapeutic response across experimental groups. **(f)** Representative confocal images of the treated and non-treated GS401-BTB assembloids over the 9-day treatment period. Scale bar, 100 μm. **(g)** Quantification plot showing the fold change of GSC counts in the AAV.CPP.16-GFAP-TK1 treated and non-treatment groups. Data are represented as mean ± s.d., from n=4 biological replicates, n_assembloids_= 8. The *P* values were calculated using two-way ANOVA with Sidak’s multiple comparison test. **(h)** Representative confocal images of the TMZ-treated and non-treated GS184-BTB assembloids over the 9-day treatment period. Scale bar, 100 μm. **(i)** Quantification plot showing the fold change of GFP⁺ GSCs counts in the TMZ treated and non-treatment groups. Data are represented as mean ± s.d., from n=4 biological replicates, n_assembloids_= 8. The *P* values were calculated using two-way ANOVA with Sidak’s multiple comparison test. **(j)** Representative confocal images of the Dox-treated and non-treated GBM-X6-BTB assembloids over 6 days. Scale bar, 100 μm. **(k)** Quantification plot showing the fold change of GFP⁺ GSCs counts in the Dox treated and non-treatment BTB assembloids and the matched neurospheres. Data are represented as mean ± s.d., from n=3 biological replicates, n_assembloids_= 4. The *P* values were calculated using two-way ANOVA with Sidak’s multiple comparison test.

To further investigate the mechanisms underlying BBB transport of BTP-7, BBB assembloids were incubated with BTP7-Cy5 under conditions that inhibit distinct endocytic and transcytotic pathways (**Extended Data Fig. 6c, d**). Robust BTP7-Cy5 accumulation was observed throughout BBB assembloids at 37 °C, whereas a Cy5-labeled control peptide (Cy5-Pas) exhibited minimal uptake, supporting the specificity of BTP-7 transport. Incubation at 4 °C markedly reduced BTP7-Cy5 fluorescence intensity, indicating that peptide uptake is energy-dependent and mediated by active cellular processes rather than passive diffusion.

Pharmacological inhibition studies further demonstrated that blocking clathrin-mediated endocytosis using chlorpromazine (CPZ) did not significantly alter BTP7-Cy5 uptake. In contrast, disruption of caveolae/lipid raft-associated transport with nystatin (Nys) significantly reduced BTP7-Cy5 accumulation within BBB assembloids. Similarly, inhibition of actin polymerization using cytochalasin D (CyD) markedly decreased BTP7-Cy5 uptake (**Extended Data Fig. 6c, d**). Together, these findings suggest that BTP-7 transport across BBB assembloids primarily occurs through active caveolae/lipid raft-associated and actin-dependent endocytic or transcytotic pathways rather than classical clathrin-mediated endocytosis.

#### Gene therapy using Adeno-Associated Virus – AAV.CPP.16

Next, we evaluated therapeutic response using AAV.CPP.16, an AAV vector previously shown to penetrate the BBB and demonstrate efficacy against GBM in vivo^31^. AAV.CPP.16 penetration was observed in both the BBB and BTB assembloids by confocal imaging, with AAV-transduced cells detected more than 50 μm below the assembloid surface in all groups. (**Supplementary Fig. 4g**).

To investigate GBM-targeting, we used AAV.CPP.16-GFAP, in which a GFAP promoter restricts transduction only to GFAP-positive cells. Western blot showed GFAP is expressed abundantly in the GSCs (**Extended Data Fig. 6e**). In the BTB assembloids, AAV signal colocalized strongly with GFP^+^ GBM cells across all groups (**Supplementary Fig. 4h - 4j**). ROI-based analysis of the mean intensity in tumor-dense regions in the BTB assembloids confirmed significant GBM-AAV overlap (**Supplementary Fig. 4i, 4j**), indicating efficient transduction of GFAP-positive GBM cells by AAV.CPP.16-GFAP.

For therapeutic testing, we used AAV.CPP.16-GFAP to deliver the suicide gene Herpes simplex virus type 1 thymidine kinase (HSV-TK1), which metabolizes the prodrug ganciclovir (GCV) into a cytotoxic compound^31^. Upon formation, the assembloids were treated with AAV.CPP.16-GFAP-TK1, and then GCV (8 μM for 8 h, followed by 16 h incubation in fresh medium) daily for over 9 days (**Fig. 6d**). The 8 μM dose was chosen because it does not alter BBB dextran permeability (**Extended Data Fig. 6f**). Confocal imaging revealed marked tumor reduction in the AAV-treated groups, while untreated BTB assembloids exhibited ongoing GBM growth and infiltration (**Fig. 6f, Extended Data Fig. 6g, 6h**). To quantify treatment response, the BTB assembloids were dissociated into single cells on specific days during the treatment course, and the number of GBM cells relative to Day 0 was measured by microscopy (**Fig. 6e)**. In the untreated groups, significant tumor progression is observed (1.8-, 3.9-, and 2.4-fold increase in tumor cell count for GBM-X6, GS184 and GS401, respectively by Day 9). In contrast, a drastic decline in tumor cell number (95-99% by Day 9 relative to Day 0) is observed in the AAV-treated groups (**Fig. 6g**). These results align with prior *in vivo* findings showing AAV.CPP.16-GFAP-TK1 and GCV therapy led to substantial tumor volume shrinkage^31^.

#### Temozolomide (TMZ) response in BTB assembloids

Next, to determine whether the platform preserves clinically relevant, line-specific therapeutic responses, we examined TMZ, the first-line agent for GBM treatment. GS184-BTB and GS401-BTB assembloids received 300 μM TMZ starting day 2, dosed 8 h on / 16 h off daily through day 9. TMZ dose was chosen based on IC_50_ results (**Supplementary Fig. 4k).** At each time point (Day 0, 3, 6, and 9), assembloids were dissociated into single cells for quantitative cell counts.

In GS184-BTB assembloids, untreated controls showed progressive tumor expansion across all time points (Day 3: ∼1.24-fold; Day 6: ∼2.22-fold; Day 9: ∼2.63-fold relative to Day 0), while TMZ-treated assembloids showed growth arrest by Day 3 (∼1.00-fold), followed by progressive cell loss through Day 6 (∼0.62-fold) and Day 9 (∼0.40-fold relative to Day 0), confirming early-onset and sustained TMZ sensitivity (**Fig. 6h, 6i**). In contrast, GS401-BTB assembloids showed no sustained reduction in tumor cell burden at any measured time point throughout the treatment course. Untreated GS401-BTB controls expanded modestly over time (Day 3: ∼1.33-fold; Day 6: ∼1.28-fold; Day 9: ∼1.59-fold), and TMZ-treated assembloids tracked closely with untreated controls at every time point (Day 3: ∼1.27-fold; Day 6: ∼1.16-fold; Day 9: ∼1.33-fold), with no significant divergence at any stage (**Fig. 6i, Extended Data Fig. 6i**). These divergent longitudinal kinetics are consistent with the primary and recurrent origins of the GSC lines: GS184 was derived from the primary tumor and GS401 from recurrence in the same patient. Together, these findings demonstrate that the BTB assembloids preserve line-specific differential drug responses, distinguishing TMZ-sensitive from TMZ-resistant GBM phenotypes.

#### Doxorubicin (Dox) delivery in BTB assembloids

We further evaluated Doxorubicin (Dox), a potent, poorly BBB-penetrant chemotherapeutic with an IC₅₀ of 36 nM in GBM-X6 cells (**Supplementary Fig. 4l**), to assess whether the BTB microenvironment influences therapeutic response. Dox was tested at 20 nM, a sub-IC50 concentration selected to evaluate BTB-associated chemoprotection while minimizing direct cytotoxicity to the assembloid. When applied to BBB assembloids, Dox treatment did not significantly alter BBB assembloid diameter over 6 days (**Supplementary Fig. 4m**); this measurement was used to assess gross morphological stability. We next treated GBM-X6-BTB assembloids with the same concentration and compared their response to matched NS cultures (**Fig. 6j, Extended Data Fig. 6j**). Quantitative cell count analysis across all time points (Day 0, 2, 4, and 6) showed that untreated GBM-X6 NS maintained relatively stable cell numbers throughout the observation period (Day 2: ∼1.00-fold; Day 4: ∼0.97-fold; Day 6: ∼1.17-fold relative to Day 0), confirming the absence of substantial culture-related changes in cell number (**Fig. 6k**). Dox-treated GBM-X6 NS, however, showed a progressive decline in cell number (Day 2: ∼1.05-fold; Day 4: ∼0.83-fold; Day 6: ∼0.82-fold relative to Day 0), indicating a Dox-specific cytotoxic effect in the absence of a barrier (**Fig. 6k**). In contrast, GBM-X6-BTB assembloids showed comparable cell counts in both Dox-treated and untreated conditions throughout the treatment course (control: Day 2: ∼1.06-fold; Day 4: ∼1.09-fold; Day 6: ∼1.21-fold; Dox-treated: Day 2: ∼1.04-fold; Day 4: ∼1.13-fold; Day 6: ∼1.27-fold relative to Day 0), with no significant difference between groups at any time point (**Fig. 6k**). These findings demonstrate that the BTB niche confers substantial chemoprotective effects that reduce Dox-associated cytotoxicity.

To investigate whether active efflux transport contributes to BTB-associated chemoresistance, we next combined Dox treatment with the P-gp inhibitor verapamil. Dox + verapamil treatment resulted in a significant reduction in GBM-X6 tumor cell number relative to untreated BTB controls by Day 6 (**Supplementary Fig. 4n, 4o**). Importantly, dextran permeability remained unchanged between untreated and Dox + verapamil-treated BTB assembloids (**Supplementary Fig. 4p**), indicating that the reduction in tumor burden was not accompanied by overt disruption of barrier integrity. Considered together with the lack of response to Dox alone in the preceding experiment and the independent Rh123 transport studies demonstrating verapamil-sensitive P-gp activity (**Supplementary Fig. 1f**), these findings are consistent with P-gp-mediated efflux contributing to Dox chemoprotection within the BTB microenvironment.

Altogether, these results indicate that the BTB assembloids provide a barrier-aware, human multicellular platform that can report drug delivery, target engagement, and therapeutic efficacy on a rapid timescale, supporting their use as a barrier-aware preclinical platform for evaluating GBM therapeutic delivery and response.

### Immune cell integration in BTB assembloids

To investigate immune cell interactions within the GSC-NVU niche, we established two complementary immune-integrated BTB assembloids by introducing either peripheral blood mononuclear cells (PBMCs) or human brain-derived microglia (HM) into BTB assembloids. Immunostaining showed higher numbers of CD3⁺ T cells on the surface of BTB assembloids compared to control BBB assembloids, with enrichment around the GSC-enriched regions in the BTB assembloids (**Supplementary Fig. 5a, 5b**). This is consistent with human GBM, where T-cell abundance is higher than in normal brain^126^. The perivascular clustering mirrors human GBM^127^, where T cells are typically sparse overall but preferentially accumulate along vessels and at tumor interfaces rather than widespread distribution within the brain parenchyma. In the microglia-BTB model, IBA1⁺ human microglia in the assembloids clustered near tumor aggregates (**Supplementary Fig. 5c**), mimicking the peritumoral microglial positioning observed in GBM tissues in mice (**Fig. 5a**).

Together, these immune-augmented BTB models capture complementary immune-vascular behaviors. The PBMC-BTB assembloid models T-cell incorporation and localization across a compromised BTB, while the microglia-BTB assembloid recapitulates microglial recruitment to tumor-vascular niches. These findings underscore the platform’s modularity and versatility for probing cell type-specific immune interactions and for studying immune-mediated modulation of BTB permeability and tumor progression.

### Integration with iPSC-derived CNS-like endothelial cells (iPSC-CECs)

To establish a genetically adaptable platform for patient-derived GSC-neurovascular interactions, we generated BBB_iPSC_ and BTB_iPSC_ assembloids using human iPSC-derived CNS-like endothelial cells (iPSC-CECs) together with primary HA and HBVP. iPSC-CECs were confirmed to express key endothelial and BBB-specific markers, including CD31, VE-cad, ZO-1 and Claudin-5 (**Extended Data Fig. 7a**). Within 2 days, BBB_iPSC_ assembloids exhibited uniform expression of tight junction and transporter proteins (Claudin-5, VE-cad, ZO-1, P-gp and Glut1) comparable to hCMEC-based BBB assembloids (**Extended Data Fig. 7b, 7c**).

By day 5, confocal imaging revealed the emergence of organized endothelial structures characterized by co-localization of CD31, VE-cad and ZO-1, which developed into increasingly branched vascular-like networks by day 7, displaying continuous junctional Claudin-5 and VE-cad expression (**Extended Data Fig. 7d, 7f**). Quantitative analysis showed significantly higher mean fluorescence intensities of ZO-1, VE-cad and CD31 in BBB_iPSC_ assembloids vs hCMEC-based BBB assembloids, indicating enhanced endothelial junctional organization (**Extended Data Fig. 7e**).

Next, we incorporated GSCs to form BTB_iPSC_ assembloids. On day 5, GSCs invaded the NVU and established close contacts with endothelial structures (**Supplementary Fig. 5d**). By day 7, endothelial remodeling produced vessel-like structures concentrated in GSC-dense zones, as indicated by CD31 staining (**Extended Data Fig. 7g)**, consistent with local, tumor-driven angiogenic cues (e.g., VEGF and other pro-angiogenic factors) that locally stimulate endothelial proliferation and vessel sprouting. Vessels adjacent to GSC clusters exhibited reduced ZO-1 and VE-cad (**Extended Data Fig. 7g**), indicative of junctional alterations in regions proximal to GSCs, mirroring the hallmark leaky vasculature characteristic of GBM *in vivo*. Western blot analysis also confirmed ZO-1 protein expression in all GSC lines used in this study (**Supplementary Fig. 5e**), supporting the interpretation that the ZO-1 signal observed within GSC-containing regions reflects genuine protein expression rather than imaging artifact.

Collectively, these findings demonstrate that BTB_iPSC_ assembloids recapitulate key pathological attributes at the GBM-NVU interface (changes consistent with tumor-associated barrier disruption), providing a flexible and genetically tractable platform for investigating line-specific neurovascular interactions, BTB permeability and therapeutic response. The ability to substitute patient-derived iPSCs and add further NVU cell types broadens the platform’s translational potential for patient-derived GBM modeling.

## Discussion

Malignant brain tumors remain among the most challenging cancers to treat, in large part because the BBB/BTB restricts therapeutic access. Conventional patient-derived GSC neurospheres have been indispensable for studying GBM biology, but they lack direct interaction with neurovascular unit (NVU) cells and cannot report how the BTB shapes tumor behavior and drug delivery. Our BTB assembloids address this gap by embedding pre-established, low-passage patient-derived GSCs within a human NVU consisting of brain endothelial cells, pericytes and astrocytes, in a protein-rich medium that more closely resembles the brain tumor microenvironment. Transcriptomic, imaging and functional assays converge to show that, compared with matched neurospheres, GSCs in BTB assembloids rapidly adopt invasive, mesenchymal, angiogenic and therapy-resistant programs. These changes translate into increased invasion *in vitro* and more infiltrative growth *in vivo*.

Correspondingly, the endothelial population in the BTB assembloids diverges from their counterparts in BBB-only assembloids (lacking tumor cells), upregulating angiogenesis, inflammatory, EMT and hypoxia pathways characteristic of GBM vasculature. Ligand-receptor analyses, together with protein-level validation of selected components such as FN1 and CD44, identify candidate niche-associated interactions at the tumor-vascular interface. We view these interaction maps as hypothesis-generating. Although we have not yet described antagonist or genetic perturbation experiments targeting specific ligand-receptor interactions, these are key next steps to define actionable mechanisms and therapeutic targets.

Direct physical contact between tumor and NVU cells emerges as a critical determinant of BTB behaviour. Tumor-conditioned medium alone does not substantially alter BBB permeability, whereas co-assembly with GSCs produces local tight-junction disruption (e.g., fragmented ZO-1/Claudin-5), astrocyte activation and increased macromolecular leakage compared with BBB-only controls. GBM-pericyte crosstalk is known to activate processes such as vascular cooption and mimicry, as well as alters pericyte transcriptome to facilitate tumor evasion of immune response^128^. Astrocyte-GBM interactions have been implicated in changes in ECM components, chemokines, cytokines and cytoskeletal rearrangements to promote tumor progression^120^. Although the roles of pericytes and astrocytes in our BTB assembloids have yet to be thoroughly examined, it is likely that these cell populations also significantly contribute to the various gene, protein and functional alterations of the GSC in the BTB assembloid. Indeed, we observe that astrocytes tend to reorganize themselves in close proximity to the tumor cells in the BTB assembloids, mirroring peritumoral astrocyte redistribution seen *in vivo*^116,117^ and play important functions in EMT, tumor invasiveness and therapeutic resistance^43^.

GSCs in the BTB assembloids also exhibit downregulated pathways related to neuronal development. In the future, other cell types of the NVU within the tumor microenvironment, such as neurons and other immune cells can be incorporated into the assembloids to further advance the model. To begin addressing the role of immune cells, we have incorporated PBMCs and microglia into BTB assembloids and observed higher levels of CD3⁺ T cells and IBA1⁺ microglia in the BTB vs BBB assembloids, with preferential localization to the perivascular and peritumoral zones. These proof-of-concept data show that this modular platform can support immune cell localization and compartmentalization, although systematic analysis of how microglia and circulating immune subsets remodel the BTB is reserved for future work.

Our assembloid model occupies a different niche than *de novo* GBM organoids. Here, we begin with GSC cells already shown to retain key features of their parental tumors. Time-course studies show that by Day 1, key junctional proteins (e.g., ZO-1 and VE-cadherin) are detectable at the endothelial barrier and remain stable through at least Day 8 (**Supplementary Fig. 6a-6c**). In parallel, barrier permeability decreased over the same time frame, reaching a stable low-permeability plateau by Day 3 that persists through Day 8 (**Supplementary Fig. 6d**). By Day 3, assembloids already display size-selective permeability (4.4 kDa vs 155 kDa dextrans) and preserved P-gp-dependent efflux. We therefore selected Day 3 as a mechanistically mature and functionally stable stage for most analyses. Across this standardized production window, we quantified size uniformity of 96 simultaneously-generated BBB and BTB assembloids per condition and observed highly consistent diameter distributions, with low coefficients of variation (CV = 4.86-5.72%), confirming robust production reproducibility (**Supplementary Fig. 6e**). A “fusion” configuration, in which pre-formed BBB assembloids are later fused with GSC neurospheres produced similar junctional disruption and permeability increases as the standard one-step assembly, indicating that BTB breakdown is driven by GBM-intrinsic pathology (**Supplementary Fig. 6f-6i**).

An important finding of this study is that increased BTB permeability does not necessarily translate into improved therapeutic accessibility. Although BTB assembloids exhibited disrupted endothelial junctions and increased dextran permeability relative to BBB controls, active efflux transport remained functional. Consistent with sustained P-gp activity in the Rh123 transport assays, verapamil co-treatment reduced GBM-X6 tumor cell number relative to untreated BTB controls without significantly altering dextran permeability. These findings are consistent with a contribution of verapamil-sensitive, P-gp-mediated efflux to Dox chemoprotection in BTB assembloids, while distinguishing structural barrier leakiness from effective drug access. Thus, transporter-mediated efflux may remain an important contributor to therapeutic response even when barrier integrity is compromised. By enabling simultaneous assessment of barrier permeability, transporter function, and therapeutic response within the same multicellular system, the BTB assembloids provide a valuable platform for dissecting the multiple mechanisms governing drug delivery across the BTB.

The detection of RFP-positive material in GSCs is consistent with EV-associated EC-to-GSC transfer and highlights the capacity of the BTB assembloids to capture multicellular communication within the GSC-NVU niche. While the present study does not establish a causal role for EV signaling in driving BTB-associated phenotypes, the platform may provide a useful system for future mechanistic investigation of EV-mediated tumor-vascular interactions and therapeutic resistance.

The BTB assembloid was designed to prioritize a combination of capabilities that remains difficult to achieve simultaneously in existing *in vitro* models, rather than maximal anatomical reconstruction of the cerebral vasculature. Perfused microfluidic systems provide greater vascular architectural complexity and physiological flow but are generally less amenable to large-scale parallel experimentation, whereas patient-derived GBM organoid models preserve tumor heterogeneity but typically lack a functionally characterized neurovascular barrier for transport studies (**Table 1**). The present platform integrates patient-derived tumor biology, a multicellular human BBB/BTB interface, matched BBB-BTB comparisons, mechanistic perturbation, therapeutic transport and response, and experimental scalability within a single system. Accordingly, its intended utility is complementary to, rather than a replacement for, anatomically more complex vascularized or *in vivo* models.

The current BTB assembloids are non-perfused multicellular tissues, not lumenized flow-bearing microvessels. Classical TEER methods optimized for 2D Transwell assays are technically impractical to adapt to free-floating 3D tissues because of electrode access, non-uniform current paths and shunting through the medium. Instead, here, we validate barrier function using orthogonal functional readouts (size-selective dextran permeability, transporter expression and P-gp-dependent efflux). Importantly, our conclusions are grounded in tumor-induced changes in barrier properties (e.g., BTB vs. BBB) together with differential handling of benchmark compounds (e.g., BBB-penetrant vs. control non-penetrant agents), rather than on shear-dependent vascular physiology. While the relative comparison between BTB and BBB conditions remains informative in this static system, absolute transport kinetics, endothelial maturation states and barrier properties are expected to be further refined under perfused conditions incorporating physiological shear stress and continuous flow. Integrating true perfusion (e.g., via microfluidic coupling or bioprinted vascular conduits) therefore represents an important future direction to further enhance vascular maturation and transport fidelity.

To begin addressing this limitation, we evaluated BBB assembloids cultured under cyclic physiological-range hydrostatic pressure (80-120 mmHg at 0.5 Hz; continuous medium perfusion at 0.1 mL/min), a hemodynamic mechanical cue that captures aspects of vascular pressure but does not reproduce luminal blood flow or endothelial shear stress. Exposure to hydrostatic pressure reduced dextran permeability and produced more compact assembloids compared with static controls, consistent with enhanced barrier integrity (**Supplementary Fig. 6j, 6k**). Although these experiments do not fully recapitulate vascular perfusion, they provide preliminary evidence that the assembloid barrier is responsive to physiological-range mechanical stimulation. Future integration of microfluidic perfusion, hydrostatic flow control, or bioprinted vascular conduits will enable more physiologically relevant modeling of blood flow and vascular transport.

Primary cells are derived from different individuals, resulting in a chimeric but genetically heterogenous NVU. To address this, we developed a proof-of-concept iPSC-based variant, in which iPSC-derived CNS-like endothelial cells (CECs) are used to generate BBB^iPSC^ and BTB^iPSC^ assembloids. By Day 2, the BBB^iPSC^ assembloids express core BBB markers, and by Day 7, we observe spatially organized tube-like structures preferentially surrounding GSC-rich regions. Although these pre-vascular networks are not yet perfused, they demonstrate that iPSC-based vascularization is feasible within this modular platform and provide an entry point for fully isogenic assembloids in which NVU cells and GSCs share a single patient’s genetic background. Our current transcriptomic analyses rely on bulk RNA-seq, which lacks single-cell resolution and spatial context. Future single-cell and spatial transcriptomic approaches will be important to resolve rare or intermediate subpopulations and to map signaling gradients within the BTB microenvironment.

This platform is designed for scalable, plate-based, high-throughput-compatible applications. BTB/BBB assembloids are formed directly in standard ultra-low attachment multiwell plates and can also be generated on micropatterned hydrogel arrays^129^, with >98% formation success and viability >2 weeks. For transcriptomic experiments, we routinely pooled >1,000 assembloids per condition, demonstrating practical large-batch production. In our prior work^35^, we have demonstrated the BBB assembloid platform supports multi-compound permeability screens in 96-well screening formats, with robust and predictive ranking reproducibility across independent runs. Here, we extend that concept to a tumor-containing, BTB-specific context with added readouts on glioma-vascular crosstalk and therapeutic response. Compared with *de novo* GBM organoids that often lack a well-characterized barrier, the BTB assembloids provide rapid, patient-derived, barrier-aware readouts at discovery throughput. Because they are modular, they can also be combined with pre-matured GBM organoids when extended tumor maturation or lineage complexity is desired, yielding hybrid systems that pair organoid heterogeneity with a defined human neurovascular interface.

Altogether, the BTB assembloid model *1)* offers a practical, scalable and mechanistically informative model that recapitulates key features of the human BTB, *2)* reveals reciprocal reprogramming between GSCs and NVU cells, and *3)* enables barrier-informed evaluation of brain tumor therapeutics, while leaving clear avenues for future refinement in perfusion, single-cell resolution and targeted mechanistic perturbation.

---

**Extended Data Fig. 1. | (a)** Characterization of the neurovascular unit (NVU) cells with cell-specific biomarkers; GFAP for human astrocytes, PDGFRβ for HBVP, CD31 and CD105 for hCMEC. Scale bar, 100 μm. **(b)** Representative confocal images showing GSC neurospheres (NS), and the self-assembled BBB and BTB assembloids after co-culturing with NVU cells in HS or NB condition for 48 h. Scale bar, 100 μm. **(c)** Mean fluorescent intensity quantification of the expression of the stem cell markers (Nestin, SOX2, and OLIG2) in GSC neurospheres from Fig.1b. The data are represented as mean ± s.d., n_assembloids_ = 4-30. The *P* values were calculated using two-way ANOVA with Sidak’s multiple comparison test. **(d)** Flow cytometry analyses showing Nestin, SOX2, and OLIG2 expression in GSC neurospheres following 72 h-culture in HS and NB. **(e)** Proliferation rate of the GSCs in neurosphere form over 4 days in either HS or NB measured by CellTiter-Glo assay. GSCs exhibit higher proliferation rate in HS compared to NB in all GSC groups. The data are presented as mean ± s.d. from n = 3 biological replicates. The *P* values were calculated using two-way ANOVA with Tukey’s multiple comparison test, **P < 0.01, ***P < 0.001, ****P < 0.0001). **(f)** Proliferation rate of the GSCs in BTB assembloids in either HS or NB culture conditions. GSCs have higher proliferation rates in HS relative to NB across all groups. The data are presented as the mean ± s.d. from n=3 biological replicates, each n contains 8 assembloids (n_assembloids_=8). The *P* values were calculated using two-way ANOVA with Sidak’s multiple comparison test. **(g)** Top left: Representative TEM image displaying the cellular organization in GBM-X6-BTB assembloid. Scale bar, 10 μm. Magnified insets 1, 2 and 3 presenting the electron microscopy features of each NVU cell. Endothelial cells feature large, oval or irregularly shaped nucleus (blue arrows, green), containing glycogen granules (small, dense particle, indicated by green arrows), and well-developed endoplasmic reticulum (ER) (yellow arrows) (inset 1, scale bar, 2 μm; inset 4, scale bar, 500 nm); Astrocytes are distinguished by abundant glial filaments (orange arrow). The presence of glycogen granules, lysosomes and autophagic vacuoles are additional features for astrocytes identification (inset 2, scale bar, 500 nm); HBVPs are identified from their well-developed ER and Golgi bodies (white arrows) (inset 3, scale bar, 500 nm). **(h)** Representative TEM images of the GBM-X6 and GS401 neurospheres: (1) Nuclear Atypia; highly irregular and convoluted nuclear (blue), scale bar, 2 μm; (2) breakdown of cristae in mitochondria (green arrows), scale bar, 500 nm; (3) intermediate cytoplasmic filaments (orange arrows) and abundant Golgi apparatus (red arrows), as well as many vesicle. Scale bar, 500 nm.

**Extended Data Fig. 2. | (a)** Representative confocal images showing the influx of TRITC Dextran signal into the BBB and BTB assembloids. GS184-BTB assembloid displays high TRITC accumulation, particularly within the tumor area (green). Dextran intensity is quantified at 50 μm depth from the assembloid surface. Scale bar, 100 μm. **(b)** Quantification of TRITC-dextran (4.4 kDa and 155 kDa) influx in BBB and BTB assembloids. BTB assembloids displayed significantly higher permeability to 4.4 kDa dextran compared to BBB controls, while permeability to 155 kDa dextran remained minimal and not significantly different among groups. The data are presented as the mean ± s.d. from n_assembloids_=3-6. The *P* values were calculated using two-way ANOVA with uncorrected Fisher’s LSD. **(c)** Bar graph showing influx of TRITC-dextran (4.4 kDa, 10 μg/mL) into the BBB assembloids after exposure to GSC-conditioned media for 72 h. Data are presented as mean ± s.d. from n_assembloids_ = 6-8. The *P* values were calculated using two-way ANOVA with Dunnett’s multiple comparisons test. **(d)** Representative images from time lapse videos showing the dynamic EC-GSC interaction and EC recruitment towards the GSCs in the BTB assembloids (G30 and GS184). BTB assembloids were embedded into bovine type I collagen (6 mg/mL) before imaging and maintained in HS. Time point (starting at 1 h) represents time post-embedding. BTB assembloids are formed with 50 G30 cells in HS condition for 24 h. GS184-BTB assembloid is formed with 500 GS184 cells in HS. EC recruitment towards the GSCs, adopting an elongated spindle-shaped morphology is observed. Scale bar, 100 μm. Representative western blot **(e)** and Quantification **(f)** showing the expression of tight and adherens junction proteins ZO-1, VE-cad and Claudin-5 in BBB and BTB assembloids. All BTB assembloids exhibited reduced levels of junctional proteins compared to BBB controls, indicating barrier disruption associated with GSC incorporation. **(g)** Representative immunofluorescence images showing the expression of efflux pump P-gp (white) in the BTB assembloids. **(h)** Quantification of Rh123 intensity in BBB and BTB with and without the treatment of P-gp inhibitor verapamil.

**Extended Data Fig. 3. | (a)** Flow cytometry analyses showing low expression of PDGFRβ in GSCs and hCMEC compared to HA and HBVP. This allows the separation of GSCs (GS184 and GS401, PDGFRβ^−^/EGFR^+^) and EC (PDGFRβ^−^/EGFR^−^) from the HA and HBVP through FACS. High EGFR expression is used to separate the GSCs from the EC (hCMEC) population. **(b)** Principal component analysis (PCA) plot showing clustering of all samples, depicting a clear separation between each group. **(c)** Venn diagram identified 143 overlapping genes (GSC_common_) that is upregulated in both the BTB_184_ and BTB_401_ groups. **(d)** Enriched hallmark pathways of the upregulated genes in GSC_common_. **(e)** KEGG pathway analysis showing the enriched pathways in BTB_401_ compared to BTB_184._ **(f)** Gene ontology annotation displaying the enriched biological processes associated with the downregulated genes in BTB_184_ and BTB_401_ compared to their corresponding GSC neurosphere (NS_184_ and NS_401_). **(g)** Hallmark analysis showing the enriched pathways in BTB_401_ compared to BTB_184._ Gene expression (average of triplicate) analysis showing genes associated with **(h)** cell cycle and **(i)** DNA repair. **(j)** Schematic illustrating the transwell invasion assay. Tranwell inserts are pre-coated with Matrigel. BTB assembloids and GSC spheres are dissociated into single cells, and seeded onto the Matrigel. Cell invasion is recorded by time-lapse imaging. Representative images show the GS184 cells seeded at T=0 h, and invaded cells at the basolateral side at T=22 h.

**Extended Data Fig. 4. | Transcriptomic analyses of EC_BTB_ and ligand-receptor interactions. (a)** Venn diagram showing 151 overlapping genes (EC_common_) between upregulated genes in EC_BTB-184_ and EC_BTB-401_, and 117 genes between downregulated genes in EC_BTB-184_ and EC_BTB-401_. **(b)** KEGG analysis showing enriched pathways of upregulated EC_common_ genes (n=151). **(c)** GO analysis showing the biological processes involved in the downregulated genes in the EC_BTB_ compared with EC_BBB_. **(d)** Heatmap of the differential gene expression of EC_BBB_, EC _BTB-184_ and EC _BTB-401_. The gene set includes 78 genes that have been identified to be upregulated in malignant GBM vasculature compared to non-malignant samples in a published study^103^. 46 (in EC _BTB-184_) and 40 (in EC _BTB-401_) genes were upregulated in the EC_BTB_ compared to the control EC_BBB_. **(e)** GO annotation showing the enrichment in biological processes, cellular components and molecular functions of the upregulated EC_common_. **(f, g)** Graphs obtained through ICELLNET showing the contribution of each molecular category to the communication scores between the GSCs and EC^105^. Paracrine communication scores are calculated through pairing the GSC_ligands_ and EC_receptor_, as well as EC_ligands_ and GSC_receptor_. For autocrine interactions, the GSC was assigned as the central cells and paired with their own cell types as partner cells. **(h)** Confocal images showing the internalization of EVs derived from RFP-expressing EC (magenta) by the GSCs (green) in the BTB assembloids. Scale bar, 20 μm.

**Extended Data Fig. 5. | Evaluation of GSCs in orthotopic PDX mouse model of GBM. (a)** Schematic showing the sample preparation of BTB_184_, NS_184_, and BBB for establishing the orthotopic PDX xenograft model. For BTB_184_, 100 BTB assembloids are pooled together, followed by papain dissociation. The dissociated cells are resuspended in NB, replated onto a 12-well plate and cultured for 2 h. This step allows for enrichment of the GSC population in suspension (while a major population of the NVU cells attached to the plate). The cell suspension is recovered and counted, and the proportion of tumor cells (tracked with CellTracker Green) relative to other cell types was measured. The concentration of GSC is adjusted to a final concentration of 80,000 GSC per 3 μL. For NS_184_, GS184 neurospheres are dissociated into single cells through papain digestion. The GSC number is counted and adjusted to a final concentration of 80,000 GSC per 3 μL. To match the cell ratio in the BTB_184_ group, NVU monoculture cells (1:1:1) are mixed with the GS184 cell suspension in the same ratio as calculated in BTB184 assembloids. As a control, BBB assembloids (100 assembloids) are dissociated and prepared. **(b)** Representative confocal images showing immunofluorescence staining and flow cytometry analysis of human vimentin showing all cells in the BBB and GS184-BTB assembloids display high vimentin expression. Scale bar, 100 μm. **(c)** Confocal images (20x) displaying the tumor distribution at 8-week post implantation (5x refer to Fig. 5a). Magnified insets 1 and 2 show enlarged view of the initial tumor implantation site and a distant tumor site in the BTB_184_ and NS_184_ groups. BBB mice injected with dissociated BBB cells as a control show no vimentin signal, indicating the NVU cells did not propagate in the mouse brain. Whole brain sections are stained with Vimentin, GFAP and IBA1 to identify the GSCs, astrocytes, and GBM associated macrophages and microglia (GAM), respectively. Scale bar, whole brain sections: 1mm; inset: 100 μm.

**Extended Data Fig. 6. | (a)** Western blot showing expression of dg-Bcan in the GSCs (purified dg-Bcan protein as positive control). **(b)** Representative confocal images displaying the BTP-7-Cy5 (white) distribution in GBM-X6-BTB and GS401-BTB assembloids compared with BBB assembloids over a 60 h-washout period (refer to Fig. 6c). **(c)** Quantification of Cy5 fluorescence intensity within BBB assembloids under the indicated treatment conditions. Data are presented as mean ± s.d. from n_assembloids_ = 5-8 BBB assembloids per group). Statistical analysis was performed using one-way ANOVA followed by Dunnett’s multiple comparisons test. **(d)** Representative confocal images of BBB assembloids incubated with 1 μM BTP7-Cy5 or Pas-Cy5 under indicated conditions for 3 h. BBB assembloids were pretreated with endocytosis/transcytosis inhibitors for 1 h prior to BTP7-Cy5 incubation. CPZ was used to inhibit clathrin-mediated endocytosis, Nys to disrupt caveolae/lipid raft-associated transport, and CyD to inhibit actin polymerization-dependent uptake pathways. Additional assembloids were incubated with BTP7-Cy5 at 4 °C to suppress energy-dependent transport processes. Gray dashed circles indicate the boundary of the BBB assembloid. Scale bars, 100 μm. **(e)** Western blot confirming GFAP expression in the GSCs and human astrocytes. **(f)** Bar graph depicting the permeability of BBB assembloids to dextran (4.4 kDa, 10 μg/mL) following exposure to GCV for 7 days. The data are presented as mean ± s.d. from n_assembloids_ = 3. The *P* values were calculated using one-way ANOVA with Dunnett’s multiple comparisons test. Representative confocal images of the treated and non-treated **(g)** GS184-BTB assembloids and **(h)** GBM-X6-BTB assembloids over the 9-day treatment period. **(i)** Representative confocal images of the treated and non-treated GS401-BTB assembloids over the 9-day TMZ treatment period. **(j)** Representative confocal images of the Dox-treated and non-treated GBM-X6 neurospheres over 6 days. Scale bar, 100 μm.

**Extended Data Fig. 7. | Generation of iPSC-derived BBB and BTB assembloids recapitulating barrier properties and GBM-associated neovascularization. (a)** Immunofluorescence images of iPSC-CECs showing expression of endothelial and BBB markers CD31, VE-cad, ZO-1 and Claudin 5. Scale bar, 100 μm. **(b)** Representative confocal images of BBB_iPSC_ and BBB_hCMEC_ assembloids displaying the expression of Claudin-5, VE-cad, ZO-1, P-gp, and Glut1 at day 2. Scale bar, 100 μm. **(c)** Quantification of MFI for Claudin-5, VE-cad, ZO-1, P-gp, and Glut1, demonstrating comparable barrier marker expression between BBB_iPSC_ and BBB_hCMEC_ assembloids. The data are presented as mean ± s.d. from n = 3 biological replicates (n_assembloids_ = 3-7). The *P* values were calculated using unpaired t test with Welch’s correction **(d)** Confocal imaging of BBB_iPSC_ assembloids in VOM condition on day 5 showing the emergence of the microvascular structures (white arrowheads) with co-localization of ZO-1, VE-cad, and CD31. Top: scale bar, 100 μm. Bottom: scale bar, 50 μm. **(e)** Quantification of MFI for ZO-1, VE-cad, and CD31 showing significantly higher expression in BBB_iPSC_ assembloids compared to BBB_hCMEC_ controls, indicating enhanced junctional organization and endothelial remodeling. **(f)** By day 7, BBB_iPSC_ assembloids displayed vascular-like networks with continuous Claudin-5 and VE-cad staining, confirming the endothelial junctional organization. Top: scale bar, 100 μm. Bottom: scale bar, 50 μm. **(g)** Representative confocal images of BBB_iPSC_ assembloids containing GSCs (GBM-X6, GS184, and GS401) showing tumor-associated vascular remodeling on Day 7. GSCs (green) infiltrate along vascular structures (CD31, magenta) and disrupt endothelial junctions (VE-cad, yellow; ZO-1, cyan). White arrowheads highlight local loss of tight junction integrity near tumor-vascular interfaces. Scale bar, 100 μm.

## Notes

### Competing Interest Statement

The authors have declared no competing interest.

### Summary of Updates

New results have been revised and updated.

